# Dynamics of drug response in single mycobacterial cells by microfluidic dose-response assay

**DOI:** 10.1101/2022.04.03.486879

**Authors:** Maxime Mistretta, Nicolas Gangneux, Giulia Manina

**Affiliations:** Institut Pasteur, Université de Paris, Microbial Individuality and Infection Group, F-75015 Paris, France

## Abstract

Preclinical analysis of drug efficacy is critical for drug development. However, conventional bulk-cell assays statically assess the mean population behavior, lacking resolution on drug-escaping cells. Inaccurate estimation of efficacy can lead to overestimation of compounds, whose efficacy will not be confirmed in the clinic, or lead to rejection of valuable candidates. Time-lapse microfluidic microscopy is a powerful approach to characterize drugs at high spatiotemporal resolution, but hard to apply on a large scale. Here we report the development of a microfluidic platform based on a pneumatic operating principle, which is scalable and compatible with long-term live-cell imaging and with simultaneous analysis of different drug concentrations. We tested the platform with mycobacterial cells, including the tubercular pathogen, providing the first proof of concept of a single-cell dose-response assay. This dynamic in-vitro model will prove useful to probe the fate of drug-stressed cells, providing improved predictions of drug efficacy in the clinic.

## Introduction

Persistence of bacterial pathogens results in long-term host colonization, which can be silent or symptomatic (*1, 2*). Persistent infections are also typically recalcitrant to treatment, due to diverse bacterial drug-escape mechanisms, with consequences for morbidity and mortality (*3, 4, 5*). Not surprisingly, pathogens responsible for persistent infections have been often implicated in drug resistance, and populate the priority list of antibiotic-resistant pathogens (*6*). Drug resistance results from spontaneous mutations, which are primarily associated with drug-target modification, drug inactivation or decreased membrane permeability, causing an increase in the minimum inhibitory concentration (MIC); conversely, drug persistence is a transient phenotypic change that is not genetically inherited, and does not cause MIC change at the population level (*4*). Drug persistence is insidious and can result from several host and pathogen factors, such as: drug absorption, distribution, metabolism and elimination (*7–9*); drug-refractory barriers built by bacterial populations, growing in the form of cellular aggregates or organized communities (*3, 10*); and phenotypic variation, an inherent feature of any clonal cell population that increases during adverse conditions. This phenotypic diversification is likely to enhance the adaptive potential of the bacterial population, contributing to drug-escape mechanisms and ultimately altering the efficacy of drugs (*11–17*). Drug persisters can be thought of as phenotypic variants within a clonal population that endure drug cidality, despite being genetically indistinguishable from drug-susceptible bacteria. Drug persistence is commonly regarded as a specific subset of drug tolerance, leading to a slowdown in the rate of drug-dependent killing, with no change in the MIC, often but not always associated with decreased growth rate (*4, 13, 18, 19*). A few studies have also begun to demonstrate a relationship between drug tolerance and resistance (*20–25*), implying that hampering drug tolerance mechanisms might prevent the onset of drug resistance.

*Mycobacterium tuberculosis* is the epitome of a persistent pathogen, and tuberculosis treatment is exceptionally protracted, due to both pharmacokinetic (PK) and pharmacodynamic (PD) limitations (*9, 26*). In particular, the tubercular pathogen occupies host niches that are refractory to drugs, where it exhibits marked growth and metabolic variation, which is associated with drug persistence (*7, 8, 11, 15*). Although non-growing bacilli are enriched with drug persisters (*1–5*), phenotypic drug-tolerance is a more complex phenomenon, contingent on variation in different cellular processes, and reported also in replicating cells (*3, 11, 13, 27*). Understanding of this phenomenon is still incomplete, mostly due to indirect investigative approaches, and this undermines preclinical drug discovery and optimization. Conventional dose-response relationship studies of drug efficacy depend on the readout used to determine the impact of drugs on pathogen fitness (*26, 28*). These approaches primarily rely on the determination of colony-forming units and of other viability parameters under static environmental conditions, inherently overlooking minor subpopulations involved in persistence (*29, 30*). In an effort to capture drug persistence, a few bulk-cell assays have been introduced (*31*). For instance, drug-dependent killing kinetics that show a biphasic pattern with a reduction in the minimum duration of killing, and growth resumption of colonies following disk-diffusion test are both suggestive of persistence. Although valuable, these approaches are limited and retrospective, especially for slow-growing pathogens such as *M. tuberculosis*, where the long time required for colony formation is likely to bring inferential and inaccurate results. As a consequence, preclinical in-vitro characterization of anti-tubercular drugs has seldom been a good predictor of sterilizing activity in humans, especially against drug-persistent subpopulations (*26, 32*). Given the complexity of bacterial drug-evasion mechanisms and their implication in therapeutic failures, preclinical study of drug efficacy requires the integration of cutting-edge approaches, which examine the PK-PD properties at higher spatial and temporal resolution, providing direct insights into the single-cell responses to drugs. Improved understanding of the dynamics of formation of drug-recalcitrant subpopulations would support decision making on the ranking of clinically relevant drugs, and provide more robust information to guide the transition from preclinical evaluation to clinical implementation (33).

Over the past two decades, nano- and micro-technologies, such as lithography, molding and device packaging, have proven their utility in biotechnology and biomedical fields (*34, 35*). The development of microfluidics-based microsystems in conjunction with time-resolved fluorescence microscopy of live cells has made it possible to quantitatively access the complexity and dynamics of microbial populations under tight environmental control with unprecedented accuracy (*36–45*). The transition from bulk-scale analysis of non-adherent cell suspensions to two-dimensional tracking of individual cells over time at the micrometric scale required enclosure systems compatible with cell size, growth, viability, changing environment and imaging. To this aim, several microfluidic systems based on different operating principles have been introduced, typically made of micropatterned polydimethylsiloxane (PDMS) and glass (*46, 47*). The most popular examples are the mother machine and its derivative microfluidic chemostats (*48, 38*). They consist of a series of narrow PDMS microchannels, whereby single progenitor cells give rise to successive generations that can be monitored until they flow into perpendicular channels, which function as both nutrient source and exit. Other PDMS-glass devices consist of growth microchannels or microchambers, of diverse shape and size, which are interconnected in fluidic networks (*42, 45, 49, 50*). While these systems are relatively functional for large adherent cells, bacterial cells find room to move, leading to three-dimensional growth and making cell tracking unstable over time. To overcome these limitations, other systems were developed to force monolayer growth, either by embedding bacteria within hydrocolloid layers, or by applying gentle mechanical compression, and providing nutrients from PDMS microchannels, which are separated from the bacterial monolayer by semipermeable membranes (*36, 37, 13*). Although these systems provide superior resolution and more stable two-dimensional growth over time, the growth areas cannot be easily multiplied nor can they be sealed independently, making these systems useless for multicondition applications. Another way to confine cells is based on the use of pneumatic valves, originally created to regulate fluids, by opening and closing multilayer networks of microchannels (*51*). Although easy to control, valve-based microsystems operate at very high pressures, which are appropriate for high-resolution imaging of intracellular compartments, but are not compatible with long-term cell viability (*39*). On the other hand, valve arrays controlled by pneumatic channels were also used to create large dome-shaped microchambers suitable either for imaging large adherent cells or for enzymatic assays on single bacterial lysates (*52, 53*). A separate category of microfluidic devices, compatible not only with cell survival and molecular analysis but also with high throughput assays, is the one based on hydrolipidic droplets (*54, 55*). Droplet-based microfluidics are being used to trap different cell types, which grow in the form of aggregates. This growth configuration impairs single-cell resolution, and requires the acquisition of several image stacks, causing phototoxicity and challenging image-analysis workflows. In summary, although several microsystems have been introduced to probe single cells, there is yet no system that stably traps very small microorganisms under separate environmental conditions, preserving their viability during prolonged periods of observation by microscopy.

Here, we report the development of a prototype microfluidic system consisting of multiple micropatterned PDMS layers bonded to glass, where an array of independent culture chambers is combined with a classical dilution tree (*56*). The peculiarity of this platform lies in its fundamental unit, which is a microchamber that traps cells through an unconventional hydro-pneumatic mechanism, allowing them to grow as a monolayer, receiving nutrients by active diffusion from the periphery of the cell-growth area. Our system is compatible with time-resolved imaging of single bacterial cells and microcolonies on a single focal plane, thus reducing phototoxicity. Furthermore, thanks to the multiplication of its fundamental unit and the integration of a double-input dilution tree, we achieve the formation of a concentration gradient, which simultaneously reaches separate chambers. The whole system is driven by a multi-channel pressure controller, allowing constant adjustment of either pressure or flow in the functional layers of the microfluidic device. With this platform, we carried out long-term imaging of both fast- and slow-growing mycobacterial cells, notably establishing a procedure compatible with biosafety level 3 (BSL3) containment. In this work, we also demonstrate the use of our system for dynamic dose-response relationship studies, by simulating PK parameters in vitro. As a proof of concept, we analyzed the relationship between concentration and effect of a relevant antitubercular drug over time, at the subpopulation and single-cell level. We found that moxifloxacin impacts mycobacterial growth in a dose- and time-dependent manner, and that clonal cells exhibit heterogeneous responses at the same drug concentration. Furthermore, while inhibition of the target occurred at higher drug concentrations, for concentrations close to the MIC the target was upregulated. We speculate that this may trigger adaptive mechanisms, leading to persistence in a fraction of the population.

We expect our microsystem to ultimately provide superior single-cell PK-PD data, which will better correlate with in-vivo efficacy. This will allow for improved predictions of drug activity against persistent subpopulations, which are challenging to eradicate, thus enhancing the control of infections.

## Results

### Construction of a microfluidic culture chamber for time-lapse imaging of bacterial cells

We conceived an unconventional microfluidic cell-culture chamber (Fig. 1A), inspired by micromechanical push-down valves (*51*), but surpassing their standards of fabrication. In particular, we designed a hydro-pneumatic microfluidic valve composed of two main PDMS layers, covalently bonded to a glass coverslip. Both the upper and the lower layer, referred to as control layer (CL) and flow layer (FL), respectively, are micropatterned with circular chambers of 1-mm diameter that are superimposed. On each layer the chambers are crossed by 300-μm wide microchannels, which are perpendicular to each other (Fig. 1A). The total height of the CL is 8 mm and the height of its micropatterned structures is 200 μm. The total height of the FL is 50 μm and the height of its micropatterned structures is about 30 μm, with a flexible PDMS top layer of about 20 μm, also referred to as membrane, whose controlled movement constitutes the operating principle of our culture chamber (Fig. 1B).

**Fig. 1.**
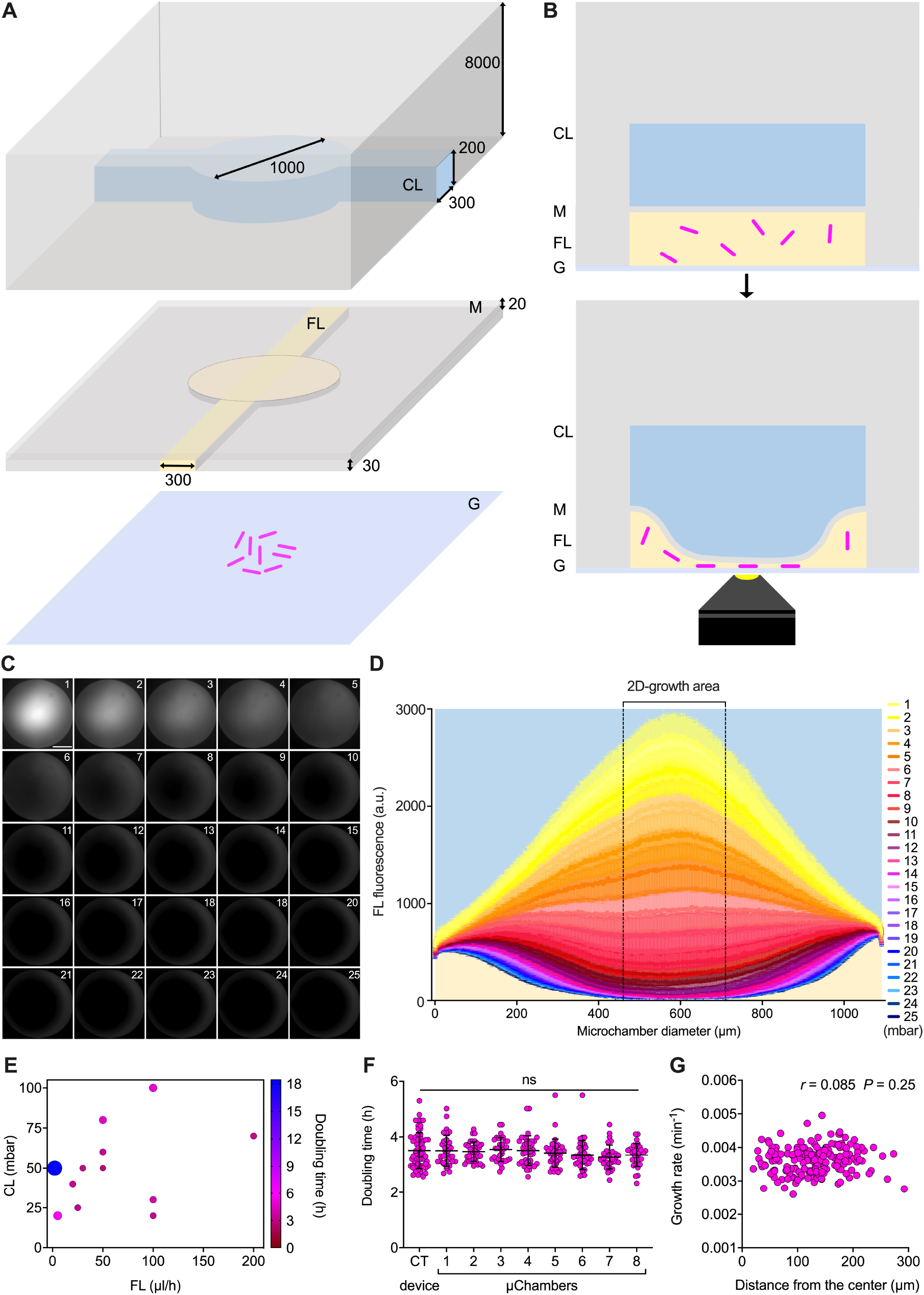
Operating principle and validation of the microfluidic culture chamber. (**A** and **B**), Schematics of the microfluidic culture chamber. Axonometric projection (A) and cross section (B) of the different layers constituting the chamber. From top to bottom are the control layer (CL), the membrane (M) integrated on the top of the flow layer (FL), and the glass coverslip (G). Magenta rods represent bacteria being trapped between M and G as the pressure in the CL increases. Bacterial growth is ensured by constant flow of the culture medium in the FL, and is monitored through an inverted microscope. Numbers represent the dimensions of the device in μm. (**C**) Representative fluorescence image series taken from the bottom of the microfluidic chamber as the pressure in the CL increases, while a FITC solution (100 μM) is injected at a constant flow rate of 150 μL/h. Numbers represent incremental pressure steps of one mbar per image in the CL, starting from 55 mbar set in the flow controller. Scale bar, 285 μm. (**D**) Fluorescence profile along the diameter of the microfluidic chamber at a constant flow rate of 150 μL/h in the FL and increasing the pressure in the CL by one mbar at a time, as in C. Fluorescence is expressed as mean ± SEM over 1690 sections (*N* = 16 chambers, from two independent experiments). CL and FL are color-coded as in A and B. (**E**) Bubble plot of *M. smegmatis* doubling time as a function of the flow rate in the FL and pressure in the CL (*N* = 24 microcolonies, from two independent experiments). (**F**) Comparison of *M. smegmatis* microcolony generation time in eight culture chambers from the same device, compared to our control (CT) device (*44*). Black lines indicate mean ± SD (33 ≤ *N* ≤ 39, from three independent experiments). No significant difference is found between microcolonies growing in independent chambers, by one-way ANOVA. (**G**) Pearson correlation between *M. smegmatis* microcolony grow rate as a function of its position in the culture chamber. The absence of correlation suggests that bacteria behave similarly at different locations within the two-dimensional growth area.

The top and bottom layers serve different but complementary functions. On the one hand, the CL is filled with water and closed at one end. By using a flow controller to fine-tune the pressure inside the CL, the PDMS membrane is lowered until it contacts the underlying glass coverslip. Lowering of the membrane creates a contact zone between PDMS and glass, suitable for microbial cell containment and two-dimensional growth (Fig. 1B). Using water instead of air to actuate the CL improves the imaging quality, since the refractive indexes of water and PDMS are similar. On the other hand, the FL is continuously perfused with growth medium, providing nutrients to the cells trapped between the PDMS membrane and the glass coverslip. Importantly, the CL is composed of a standard (1:5) ratio between PDMS crosslinker and pre-polymer, as opposed to the FL, composed of a (1:20) ratio, in order to bond the two layers by migration of the crosslinking agent (57). In addition, we found that treating the FL with oxygen plasma improves the penetration of liquid culture medium (see also Materials and Methods), due to increased hydrophilicity (*58*).

We fabricated a first microsystem prototype, by arraying eight culture chambers in a row, fed by a common flow and controlled by a single CL, laid out perpendicular to the FL (Fig. S1A). To prevent accumulation of debris in the FL and clogging of the system, we also integrated PDMS pillars of 100- and 20-μm diameter at the inlet port of the FL. To test the membrane motion, we perfused the FL with a solution of fluorescein isothiocyanate (FITC) at a constant rate of 150 μL/h, measured at the outlet, bringing the membrane to a convex position relative to the coverslip. Next, we applied a gradient of pressures to the CL, and acquired sequential images of the chamber at increasing pressure steps of one mbar per image (Fig. 1C). By measuring the fluorescence profile along the membrane diameter, we found that movement of the membrane from convex, to flat, to concave, is associated with a decrease in fluorescence (Fig. 1D and movie S1). Starting from the point of maximum convexity of the membrane, an increment of 5 mbar was sufficient to flatten the membrane, and a further increment of 20 mbar generated a contact zone of approximately 250 μm in diameter (Figs. 1D and S1B). This contact area is sufficiently large to visualize the growth of several bacterial microcolonies, and leaves sufficient room for lateral flow of growth medium. The unusual aspect ratio between diameter and height of the lower microchamber (33: 1), over three fold above the manufacturing standards typically used to prevent collapse of microstructures (*51*), enabled us to use actuation pressures in the CL more than one order of magnitude lower than those used in conventional micromechanical valves (*59*). Our ability to work at low pressure in the CL was likely to be beneficial to cellular viability.

Next, we used the FL to load a suspension of *Mycobacterium smegmatis* cells, a non-pathogenic tuberculosis model that duplicates in vitro approximately every 3 hours. To identify the best culture conditions, we tested different combinations of pressure and flow rate in the CL and in the FL, and measured the doubling time of microcolonies within the two-dimensional growth area (Fig. 1E). We found that several combinations are compatible with bacterial survival and normal doubling time. However, a drastic decrease of flow rate in the FL impaired bacterial growth. This was likely due to decreased nutrients influx and possibly to excessive pressure exerted on the cells from the CL. Based on these results, we were able to choose the operating conditions of the device that are most compatible with mycobacterial growth, namely, high flow rate in the FL (150 μL/h) and moderate pressure in the CL (30 mbar). Under these conditions, we not only measured similar generation times between separate microfluidic culture chambers (Fig. 1F), implying good reproducibility across the system, but also found that mycobacterial growth was comparable to that obtained with our former single-condition microfluidic device, which is based on a different working principle (*13, 44*). In addition, we calculated that the growth rate of microcolonies was independent of their location in the chamber (Figs. 1G and S1B), suggesting homogeneous growth throughout the two-dimensional growth area.

In conclusion, we constructed a modular microfluidic culture chamber based on a hydropneumatic mechanism. We also proved that this device can trap live bacterial cells, and can be used to monitor their two-dimensional growth.

### Evolution of the microfluidic culture chamber into a multi-condition platform

To increase the versatility of our microfluidic culture chamber, we arrayed twenty of them, generating five separate groups of four microchambers (Fig. 2A, Data S1), and we connected each group to the outlet branch of a classic microfluidic dilution tree (*56, 60, 61*). In this configuration, the CL consists of a straight microchannel that widens in correspondence of each microchamber, ensuring the same pneumatic mechanism of the single functional unit (Fig. 1). The resulting five-condition platform is supervised by a computer-driven multichannel flow-controller. The first channel regulates a tripartite tubing, by applying pressure into three separate bottles, containing either culture medium alone or the highest concentration of the molecule to be tested dissolved in culture medium (Figs. 2A and S2A). In addition, a bidirectional switch valve is placed downstream of two bottles to switch between conditions. The second channel regulates a water supply for the actuation of the CL. The third channel is connected to a sealed waste receptacle, and used to maintain a stable flow rate inside the FL, in association with a flowmeter placed downstream of the outlet port (Fig. 2A).

**Fig. 2.**
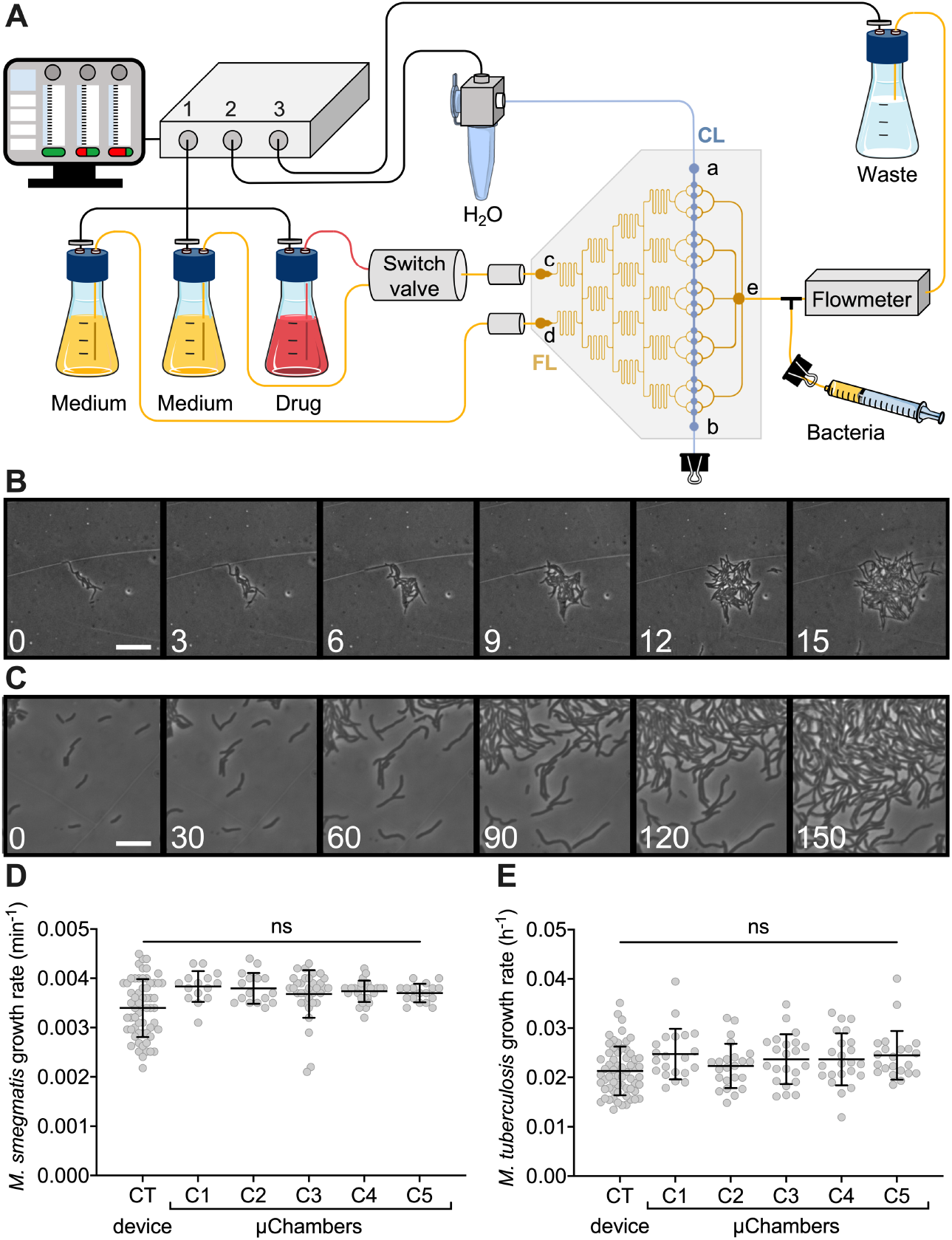
Five-condition platform set-up and validation. (**A**) Schematic of the five-condition microfluidic device connected to a multi-channel microfluidic flow controller (Fluigent). The first channel is connected to three independent reservoirs, in turn connected to a software-controlled bidirectional position valve to switch between conditions. The reservoirs are connected to two inlet ports (c and d) of the FL (yellow pattern), and two capacitors (gray cylinders) are integrated upstream of the inlet ports. A software-driven flowmeter is integrated downstream the outlet port of the FL (e), which flows into a sealed trash. The outlet tubing is bifurcated via a Y-connector, for bacteria injection. The second channel is connected to the water reservoir, which actuates the membrane from the CL (blue pattern). Filters (0.2 μm) are used at the entry of reservoir and waste bottles to prevent contamination of both fluidics and environment. (**B** and **C**) Representative phase-contrast time-lapse image series of exponentially growing *M. smegmatis* (B) and *M. tuberculosis* (C) inside the five-condition device. Cells were imaged at 20-min (B) or 3-h intervals (C), and numbers represent hours. Scale bars, 10 μm. (**D** and **E**) Microcolony growth rates of *M. smegmatis* (D) and *M. tuberculosis* (E) growing inside different chambers of the five-condition device, and compared to our CT device (*44*). Black lines indicate mean ± SD (15 ≤ *N* ≤ 60, from at least two independent experiments per strain). No significant difference is found by one-way ANOVA followed by Kruskal-Wallis multiple comparisons test.

We tested whether the five-condition platform is suitable for monolayer growth of both fast-growing *M. smegmatis* and slow-growing *Mycobacterium tuberculosis*. To this aim, we established a specific sequence of actions to prepare, assemble and load the different components of our platform, complying with the standards for BSL3 containment (Figs. S2A-C). The FL was loaded with the bacterial suspension from a secondary outlet tubing. Following a short period of incubation, the control layer was actuated to block bacterial cells between the PDMS membrane and the coverslip (Fig. S2B). After checking for leaks, the reservoir and waste bottles were connected to the device and to the flow controller, and the whole platform was mounted on the microscope stage (Figs. 2A, S2C, S3A and B). By carrying out time lapse microscopy (Figs 2B, 2C and S3B), we not only confirmed that both mycobacterial species grew at standard growth rate within different microchambers of the five-condition platform, but also proved its stability for long-term imaging (Figs. 2D and E).

### Formation of different gradient patterns

To generate a gradient of concentrations in our five-condition platform (Fig. 2A), we inject two different solutions from the inlet ports, reaching two independent serpentine-shaped microchannels. The latter bifurcate into an outer serpentine, where the solutions remain at their original concentration, and an inner serpentine that acts as a mixer, where the incoming solutions are blended. Each of the three serpentines bifurcates, originating two outer serpentines and two mixers, which further bifurcate and give rise to the last dilution level with two outer serpentines and three inner mixers. The flow in the serpentines is the same at each dilution level, while the flow in the connecting sections at each bifurcation is greater towards the outside of the system and smaller towards the inside (*60*). The concentration gradient resulting from this network structure is linear. From the last dilution level, the solutions flow into five separate clusters (C1 to C5) of four microchambers each. Finally, each group of chambers is connected to the same outlet port, aiming to maximize flow stability in the whole microsystem. We illustrate the operation of our platform by sequential mixing of dyes (Fig. 3A).

**Fig. 3.**
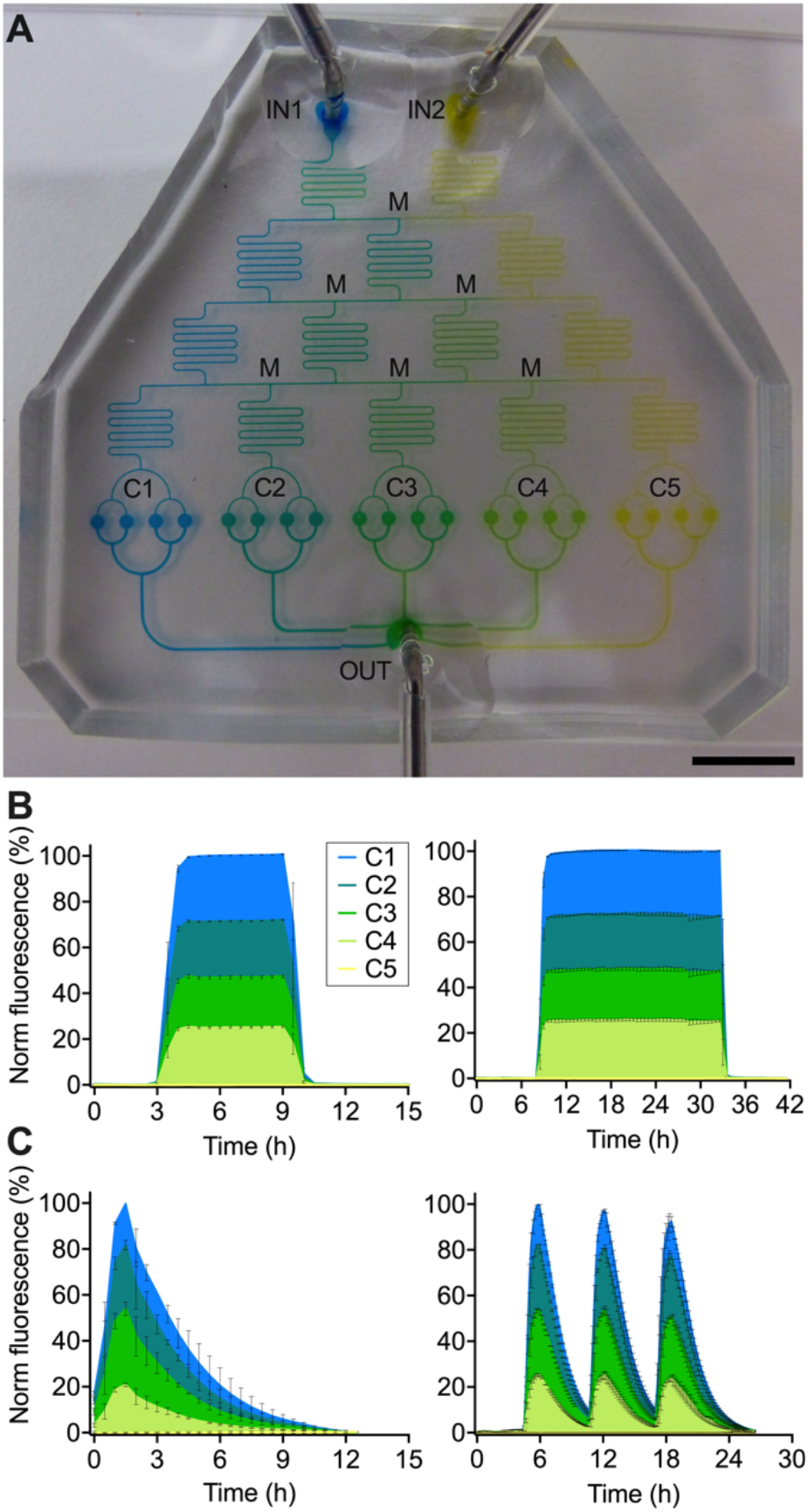
Characterization of gradient formation in the five-condition device. (**A**) Picture of the five-condition device filled with dyes from two separate inlet ports (IN1 and IN2) of the FL, in the absence of CL. Blue and yellow dyes form a color gradient in the dilution tree, via serpentine mixers (M), reach five independent groups of four chambers (C1 to C5), and flow into a common outlet (OUT). Scale bar, 10 mm. (**B** and **C**) Formation of gradients through progressive mixing of a FITC solution (100 μM) injected into IN1 with a non-fluorescent solution injected into IN2. Mean fluorescence is measured from each culture chamber. Steady (B) and pulsing (C) gradient patterns of different duration are shown. Filled lines and error bars indicate mean ± SEM (*N* = 2 or 4). Colors indicate different groups of chambers, color coded as in A.

We quantified the gradient by injecting a solution of FITC dissolved into 7H9 medium into the first inlet port, and non-fluorescent 7H9 medium into the second inlet port (Fig. S4A). At steady state, the measured gradient minimally differed from the theoretical expectations (*60*), in that the FITC solution was undiluted in C1, diluted to about 72% in C2, 48% in C3, 25% in C4, and was absent in C5 (Figs. 3B and S4A). Aiming to mimic a typical PK profile (Fig. S4B), we also generated pulse-shaped FITC gradients (Fig. 3C), whose concentrations at peak differed from those obtained with static gradients. In particular, the FITC solution at the time of maximum concentration was undiluted in C1, diluted to about 82% in C2, 54% in C3, 21% in C4, and was absent in C5. Static and dynamic gradients (Figs. 3B and 3C) were produced by alternating the perfusion of non-fluorescent and fluorescent solutions for different time intervals, by means of a bidirectional switch valve. Interestingly, we could obtain pulsing PK-mimetic gradients only when capacitors were integrated upstream of the inlet ports (Figs. 2A and S2C). In summary, by integrating a dilution-tree network with our microfluidic culture chambers, we created a microfluidic platform suitable to titrate molecules over space and time.

### Time-resolved imaging of *M. smegmatis* stressed with a pulsing gradient of moxifloxacin

To test the operation of our platform, we examined the effect of different drug concentrations on the growth rate of *M. smegmatis*. We focused on moxifloxacin, a fluoroquinolone drug targeting the type II topoisomerase DNA gyrase, and critical for antitubercular therapy against multidrug-resistant strains (*33*). To monitor the effect on the target, we generated a red fluorescent reporter of GyrA, by fusing the *mCherry* gene in frame with the *gyrA* chromosomal copy (Fig. S5A). We seeded log-phase cells of *M. smegmatis* GyrA-mCherry reporter inside the 5-condition platform (Fig. 2A). First, we grew the bacteria under constant perfusion of fresh 7H9 growth medium for six hours (about two generation times), next we injected moxifloxacin from one inlet port to generate a pulsing gradient, and finally we switched back to drug-free medium (Fig. 4A and movie S2). To mimic the PK profile of moxifloxacin (*62*), we empirically formed the gradient, producing a positively skewed injection pattern (Figs. 4B and S4B), and scaling its duration to the generation time of *M. smegmatis*. At one end of the device we left a group of chambers without drug to monitor the normal growth, and at the opposite end we tested the highest drug concentration (250 ng/ml; 5X-MIC), reaching the peak of the pulse about one and a half hour after injection (Fig. 4B). The concentrations of moxifloxacin present in the three middle groups of microchambers were estimated from the dilution factors measured by fluorescence (Fig. 3C).

**Fig. 4.**
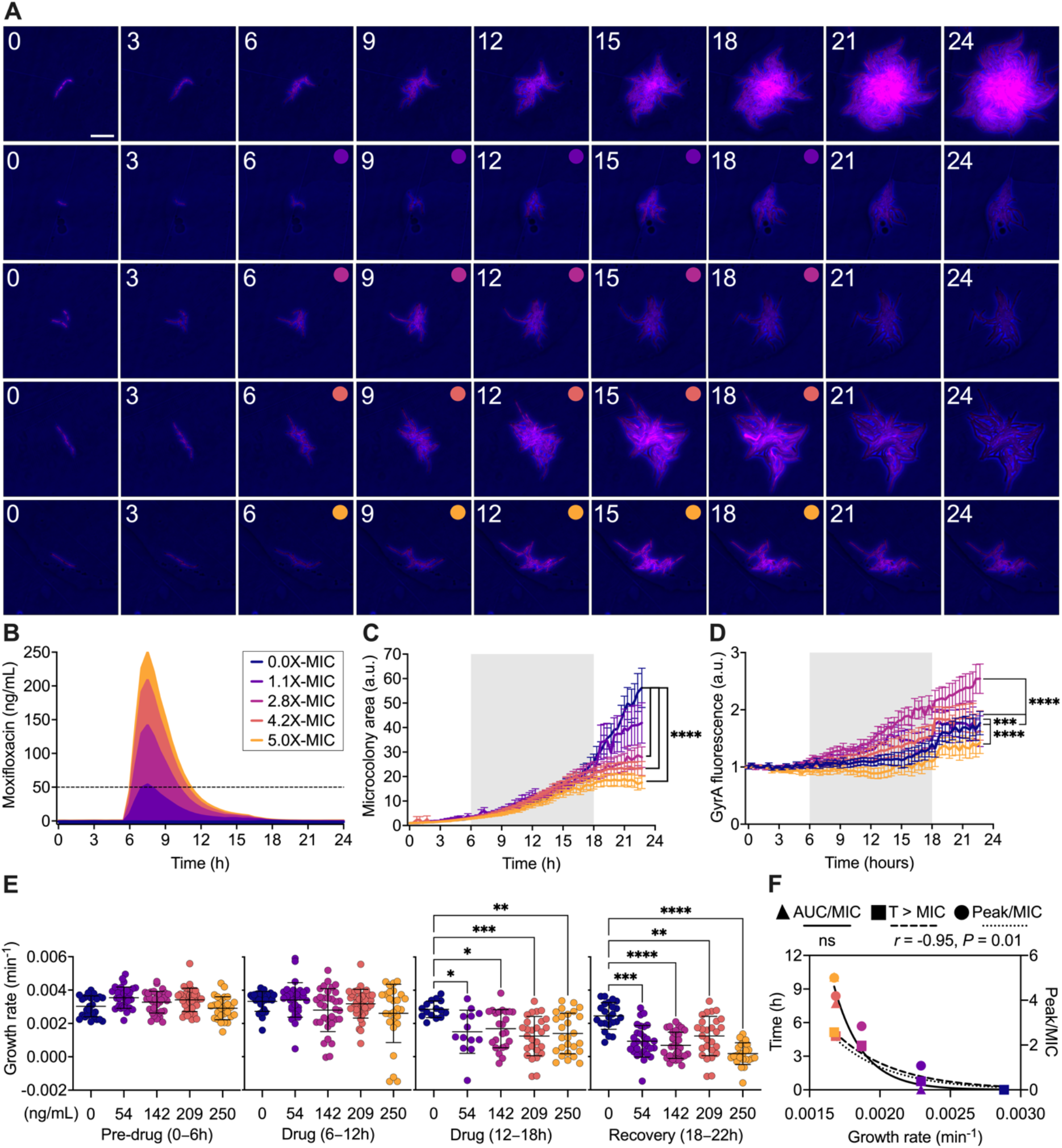
Effect of a pulsing moxifloxacin gradient on *M. smegmatis* GyrA-mCherry reporter. (**A** and **B**) Time-lapse microscopy of exponentially growing *M. smegmatis* seeded into the five-condition device and stressed with a pulsing gradient of moxifloxacin. Time-lapse image series representative of each chamber group (A). Phase-contrast and fluorescence images were acquired every 20 min and numbers represent hours. Scale bar, 10 μm. Absence of drug is unlabeled and circles indicate the presence of different drug concentrations, color-coded in (B). Concentrations shown in the inset relative to the MIC (dashed line) refer to the peak concentration reached during moxifloxacin exposure, according to the dilution factors measured in Fig. 3C. (**C** and **D**) Normalized microcolony area (C) and GyrA-mCherry fluorescence (D). Gray shading indicates drug exposure period. Lines and error bars are color-coded as in (B) and represent mean ± SEM (10 ≤ *N* ≤ 42 microcolonies, from three independent experiments). Significance by 2-way ANOVA followed by Dunnett’s multiple comparisons test: \*\*\**P* = 0.0005; \*\*\*\**P* < 0.0001. (**E**) Microcolony growth rates measured for each condition color-coded as in (B), during different experimental phases, before (pre-drug), during (drug) and after moxifloxacin exposure (recovery). Lines and error bars represent mean ± SD (10 ≤ *N* ≤ 42). Significance by one-way ANOVA followed by Dunnett’s multiple comparisons test: \**P* = 0.01; \*\**P* = 0.001; \*\*\**P* = 0.0003; \*\*\*\**P* < 0.0001. (**F**) Pearson *r* correlation between different PK parameters and averaged microcolonies growth rates (10 ≤ *N* ≤ 42), measured over the entire duration of the experiment, for each condition color-coded as in (B). Left *y*-axis: ratio of area under the concentration-time curve to MIC (AUC/MIC), and time above MIC (T>MIC); right *y*-axis: ratio of peak concentration to MIC (Peak/MIC).

We implemented a semi-automated microcolony analysis workflow (Fig. S5B, Data S2-S7), enabling us to measure size and fluorescence of several microcolonies per condition, for the entire duration of the experiment (Figs. 4C and D). While in the absence of moxifloxacin bacteria exhibited exponential growth, we measured a significant and progressive decrease of microcolony size at increasing concentrations of drug, and a rapid growth arrest at the highest concentration (Fig. 4C). Compared to untreated cells, we observed significant target inhibition at the highest concentration of moxifloxacin, and target induction at lower concentrations (Fig. 4D), with implications for the induction of DNA damage response mechanisms (*63*) and possible consequences for the onset of mutations. During the first six hours of drug exposure, we did not measure significant changes in growth rate among microcolonies growing in the absence or presence of different drug concentrations, whereas the growth rate started to slow down significantly during the last six hours of drug exposure and continued to decrease during the recovery phase in the absence of drug (Fig. 4E). In addition, we observed marked heterogeneity in growth rate among clonal microcolonies exposed to the same drug concentration (Fig. 4E). Finally, we checked whether there was a correlation between growth rate and the PK parameters simulated in vitro (Figs. 4B and S4B). Interestingly, we measured an inverse relationship between the growth rate and both dose (R^2^ = 89.8%) and time above the MIC (R^2^ = 90.8%). In contrast, we found no significant correlation between the growth rate and the area under the curve (AUC) to MIC ratio (Fig. 4F).

To conclude, we applied our platform to analyze PD properties and PD-PK relationships at high cellular resolution in space and time, and found that moxifloxacin exerts its effect on *M. smegmatis* microcolonies in a dose- and time-dependent manner.

### Single-cell dose-response assay in *M. tuberculosis* stressed with a static gradient of moxifloxacin

To test the stability of our platform for long-term imaging, we examined the effect of different concentrations of moxifloxacin against *M. tuberculosis* by multi-phasic time-lapse microscopy. We created the red fluorescent reporter strain of GyrA in *M. tuberculosis* (Fig. S6A), and further labeled it with a constitutively expressed green fluorescent protein (GFP_cyt_). While GyrA-mCherry provided information about the moxifloxacin target, the cytosolic marker helped us to identify individual bacilli and to recognize their loss of viability or lysis from gradual or abrupt fluorescence decrease, respectively. We first seeded log-phase GFP_cyt__GyrA-mCherry bacilli inside the 5-condition platform, and grew them in fresh medium for almost four days (about four generation times). Next, we injected the maximum moxifloxacin concentration (250 ng/ml; 5X-MIC) from one inlet port to generate a stable gradient for two days, leaving the opposite end of the device devoid of drug. Finally, we switched back to drug-free medium for five days to monitor the recovery phase (Fig. S6B and movie S3). Also in this case, the concentrations of moxifloxacin present in the three middle groups of microchambers were estimated from the dilution factors previously measured by fluorescence (Fig. 3B).

To increase the analytical resolution, we quantified individual bacilli and their fate over time. In the absence of drug, the number of bacilli grew exponentially until they saturated the field of view, leading to a decrease of the division rate and making cell counting impractical (Fig. 5A). At increasing concentrations of moxifloxacin, *M. tuberculosis* cell number plateaued, and the rate of division decreased until it stopped (Fig. 5A). Interestingly, cells subjected to lower drug concentrations exhibited significant increase in size, consistent with induction of DNA damage response (*13, 63*). Conversely, bacilli stressed with the highest concentration of moxifloxacin (250 ng/ml; 5X-MIC) underwent abrupt growth arrest without significant changes in their size (Fig. 5B). Overall, these results imply that small differences in drug concentrations, close to the MIC, could have important implications for the induction of adaptive stress responses in individual cells, and for the onset of drug resistance. As drug concentration increased, we also observed a progressive decrease in GFP_cyt_ fluorescence, concurrent with an increase in GFP_cyt_-dim bacilli, suggestive of gradual loss of cell viability (Figs. 5C-E). In contrast, the presence of residual GFP_cyt_ fluorescence in intact bacilli that do not grow may be attributed to a persistent-like state, which could eventually revert to active growth under appropriate conditions and in due time. Thus, in this experimental set-up, we regarded as survivors both cells that resumed growth after drug removal and those that retained weak GFP_cyt_ signal in the absence of growth. The fraction of these viable cells decreased considerably starting from a dose of moxifloxacin equal to 2.4X-MIC, when GFP_cyt_-dim cells were prevailing but only a few cells lysed (Fig. 5F). We measured a half maximal inhibitory concentration (IC50) just below the MIC value, while the maximum drug efficacy was achieved at 2.4X-MIC (Fig. 5G). This was also consistent with the effect on the target, in that only cells treated with the lowest concentration (1.25X-MIC) exhibited an increase of GyrA-mCherry, consistent with induction of DNA damage response by fluoroquinolones (*13, 63*). In contrast, GyrA-mCherry expression decreased at higher concentrations (Figs. 5H and S6C), implying dose-dependent inhibition.

**Fig. 5.**
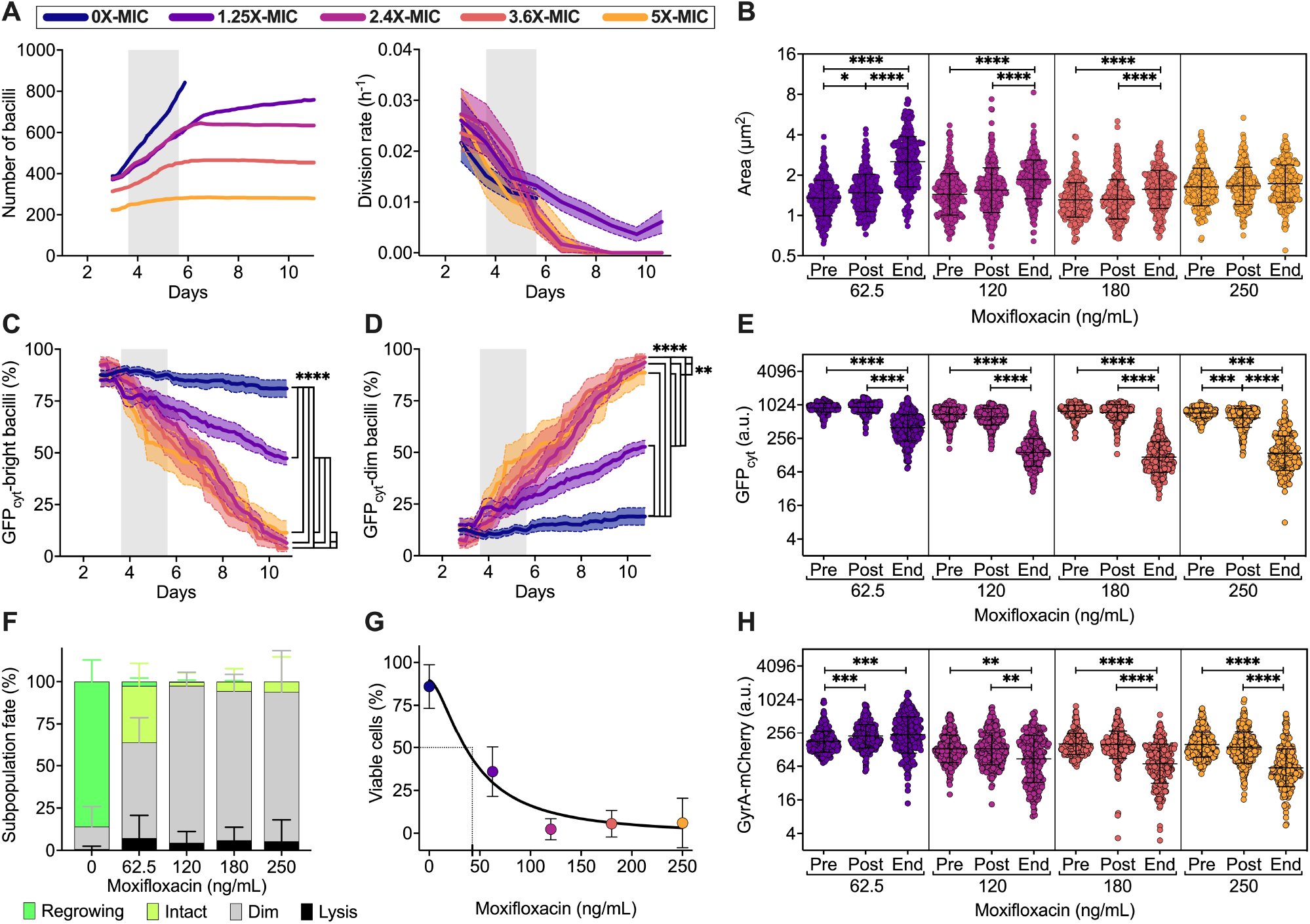
Effect of a static moxifloxacin gradient on single *M. tuberculosis* GFP_cyt__GyrA-mCherry cells. (**A**) Assessment of *M. tuberculosis* GFP_cyt__GyrA-mCherry growth during multi-phasic time-lapse microscopy. Number of bacilli (left panel) and cell division rates (right panel) in different chamber groups of the five-condition device before, during and after moxifloxacin exposure (gray shadings). Lines and error bars represent mean ± SEM (279 ≤ *N* ≤ 842 bacilli, from two independent experiments). Colors refer to the concentration of moxifloxacin used in each chambers group relative to the MIC (50 ng/mL) according to the dilution factors measured in Fig. 3B, as denoted in the inset. In the absence of drug (blue lines), cell count stops earlier due to excessive growth. (**B**, **E** and **H**) Single-cell size (B), GFP_cyt_ (E) and GyrA-mCherry (H) fluorescence at different experimental time points just before (Pre) and just after (Post) treatment, and at the end of the recovery period (End). Black lines indicate mean ± SD (253 ≤ *N* ≤ 308 bacilli, from two independent experiments). Significance by one-way ANOVA followed by Kruskal-Wallis multiple comparisons test: \**P* = 0.05; \*\**P* < 0.005; \*\*\**P* < 0.0008; \*\*\*\**P* < 0.0001. (**C** and **D**) Fractions of subpopulations over time displaying bright GFP_cyt_ (C) or dim GFP_cyt_ (D) fluorescence. Gray shadings represent the period of drug exposure. Lines and error bars represent mean ± SEM (222 ≤ *N* ≤ 1327 bacilli, from two independent experiments). Significance by two-way ANOVA followed by Tukey’s multiple comparisons test: \*\**P* = 0.006; \*\*\*\**P* < 0.0001. (**F**) Fractions of subpopulation fates at different drug concentrations (295 ≤ *N* ≤ 1327, from two independent experiments). (**G**) Cumulative fraction of survivors (regrowing and intact GFP_cyt_-bright) at the end of the recovery period as a function of moxifloxacin concentration. Symbols and error bars represent mean ± 95% CI (295 ≤ *N* ≤ 1327, from two independent experiments). Black line represents fitting of the data with a four-parameter dose-response curve: R^2^ = 0.88; d.f. 98. The IC50 value is shown (dashed lines).

Here, we report the first proof of concept of a dynamic dose-response microfluidic assay with a slow-growing and clinically relevant pathogen. We proved the functionality of the platform for longterm microscopy in BSL3, and the ability to visualize diverse single-cell responses according to small changes in drug concentration. Understanding of these dynamics is crucial for predicting adaptive responses of the pathogen in vivo under fluctuating drug concentrations. In conclusion, our platform has the potential to improve preclinical characterization of drug efficacy and the understanding of the mechanism of action. This approach can complement conventional bulk assays, offering direct information on the dynamics of killing and survival of drug-evading cells, responsible for the loss of efficacy of antimicrobial agents.

## Discussion

The ability of some pathogenic bacteria to tolerate prolonged exposure to antimicrobials can be conducive to antimicrobial resistance (*20–25*), and poses major limitations to the management of infections (*1, 64*). Although this phenomenon has been known for a long time (*3, 4*), direct identification and analysis of drug-persistent subpopulations are troublesome, hindering preclinical identification of drugs effective against them. As opposed to bulk-cell studies, spatiotemporal analysis of single bacterial cells cultured inside microfluidic devices offers insights into the dynamics of cellular responses to antimicrobial treatment, and the formation of drug-persistent cells (*11, 13, 20, 40, 43*). Despite the potential of this approach, the challenge remains of how to scale single-condition devices to multicondition platforms, while preserving device stability, user control and long-term cell viability.

Here we report the construction of a modular microfluidic cell-culture chamber that operates on a hydro-pneumatic mechanism, and is characterized by a particularly high aspect ratio. By multiplying this functional unit and combining it with a double-entry microfluidic dilution system, we derived a 5-condition microfluidic platform. The latter is composed of two inlets followed by a series of serpentine channels, which sequentially achieve titration of the incoming solutions across five independent groups of chambers. Time-resolved observation of these chambers by wide-field inverted microscopy allowed us to directly capture growth, mortality and survival rates of single mycobacterial cells before, during and after drug exposure. We tested the operation of our platform using both non-pathogenic and pathogenic mycobacterial species, which are particularly small, below 0.5 μm in thickness and between 1 and 6 μm in length, and have generation times ranging from 3 to 24 hours. Mycobacterial cells were cultured within a two-dimensional growth area, generated by gentle adhesion of a thin PDMS membrane on a coverslip. We probed single-cell growth relative to drug gradients, which were generated under diverse patterns of perfusion. Overall, this work reports the first proof of concept of a single-cell doseresponse assay for slow-growing bacterial cells, and demonstrates the ability of our platform to investigate PK-PD relationships at high spatial and temporal resolution. As opposed to bulk-cell methods that indirectly infer the presence of drug-persistent cells, our platform allows for direct detection of diverse single-cell fates over time, providing a technological advance in drug discovery.

Microfluidic devices used to trap, culture and image living cells are mostly made of micropatterned PDMS, which can be activated or functionalized on its surface, to generate irreversible chemical bonds with compatible materials, such as glass or other polymers (*46*). Typically, surface activation is obtained inside a vacuum chamber, by ionization of oxygen via an electric field that oscillates as a function of time. This so-called plasma treatment removes impurities and generates functional silanol groups, which make the surface hydrophilic, and covalently bind the same groups generated on other substrates (*65*). To seal our platforms, we use this plasma treatment followed by manual alignment, in the absence of pressure from the control layer, aiming to prevent sticking of the PDMS membrane to the underlying glass. However, the current manufacturing process of our 5-condition platform has a success rate around 80%, primarily due to manual alignment errors between the different layers that constitute the device, and to patchy adhesion of the PDMS membrane to the glass surface in some growth microchambers. These limitations in the manufacturing of the platform can impair the stability of the system, and the delivery of nutrients to cells as well as their survival. Many variables are involved in the process of surface oxidation of PDMS, thereby producing inter-device variation, which is hard to predict.

By fine-tuning plasma dose, exposure time and pressure, the surface oxidation of PDMS can lead to local vitrification (*66*). Vitrification increases the stiffness of PDMS and leads to the formation of nanometer-sized wrinkles, which could not only prevent unintended bonding, but also improve the circulation of the nutrients within the 2D-growth area. Although PDMS vitrification holds promise for device optimization, evaluation of the conditions, functioning of the platform and cell viability remain to be tested. In order to make the manufacturing process increasingly reproducible, PDMS could also be blended with amphiphilic or hydrophilic additives (*67*). The resulting hybrid materials exhibit more-stable properties over time, and are less susceptible to non-specific adsorption of hydrophobic molecules, possibly improving the circulation of drug candidates. In alternative, fabrication based on photolithography and multilayer soft-lithography could be attempted using transparent and biocompatible plastic polymers and software-controlled multilevel microfabrication techniques, such as plastic injection molding and additive stereolithography (*68*). Irrespective of the manufacturing process, the modularity and inherent versatility of our platform make it suitable to structural modifications and upgrades. For instance, it could be scaled up by increasing the number of inlet ports, branches of the dilution tree and microchambers. However, any modification from the original design will require verifying device stability, gradient formation and homogeneous cell growth across microchambers from scratch.

Since our platform was designed to trap and monitor the behavior of very small bacterial cells, using a flexible PDMS membrane that exerts low pressure on cells, allows gases to permeate and nutrients to circulate, and limits shear stress, it will most likely be compatible with a broad spectrum of microbial cells, eukaryotic cells, organoids, and may be adaptable to more complex multicellular models (*69, 70*). Additionally, the current platform allows us to combine experimental conditions in a virtually unlimited manner. For instance, nutrients, cell stimuli, flow dynamics, atmosphere and pressure can be combined and modified as a function of time, aiming to simulate different environmental micro-niches. In particular, to study the effect of drugs it is important to reproduce fundamental PK parameters (*70*). Although at present our system does not include components of the immune response, we can create diverse drug exposure patterns, and quantify the dynamics of response of individual cells under clinically relevant infusion patterns. To this aim, target-controlled infusion software systems could be implemented, in conjunction with multiple inlet ports or multi-directional valves, aiming to fine-tune each fluidic input according to in-silico models inferred from human PK parameters (*71–73*).

In-vitro models enabling accurate PK-PD analysis with high predictive potential for the in-vivo phase are essential but limited. Chief among them is the so-called hollow fiber, a well-known dynamic macrosystem used for preclinical assessment of drug activity and selection of dosing strategies, also for antitubercular drugs (*74, 75*). The hollow fiber is a cylindrical bioreactor controlled by peristaltic pumps, where the flow of nutrients and drugs reaches bacterial cells by diffusion through a series of semi-permeable capillary fibers. Growth inhibition and recovery during fluctuating drug exposure, which mimics typical human PK profiles, are monitored by regular sampling and assessment of colony-forming units (*75–77*). As a consequence, the hollow-fiber system is inherently limited by the ability of colonies to resume growth on plate, and cannot detect cells that are non-growing but metabolically active and pose a risk for relapse (*11*). Unlike the hollow fiber, our microsystem can determine PD parameters such as division and killing rates at the single-cell and microcolony level, and correlate them with critical PK parameters, also providing real-time insights into bactericidal versus bacteriostatic activity and risk of relapse for individual cells. Additionally, several drug doses can be tested simultaneously, surpassing the throughput of standard microfluidic systems (*46, 47*). Finally, using appropriate fluorescent reporters, we can also monitor the dynamics of induction and repression of the drug target as a function of drug dose and exposure time. The analysis presented here reveals not only heterogeneous responses to the same drug concentrations in individual cells, but also how small changes in drug concentration can impact the population dynamics, conceivably triggering survival mechanisms that might be conducive to drug resistance (*20–25*).

Because effective antimicrobials are inexorably running out, posing one of the biggest challenges of the 21^st^ Century (*6*), we urgently need new tools to help us better understand how drugs work and how best to use them. Our platform will prove useful as a dynamic single-cell PK-PD in-vitro model, to clarify the activity of antimicrobials and how certain bacterial cells escape treatment. Furthermore, the integration of single-cell PD datasets with clinical ones, through mathematical modeling and simulations (*78, 29*), might also help to better predict in-vivo efficacy. We ultimately hope that our work will contribute to the establishment of more robust drug-development pipelines, aiming to prevent disease relapse and to improve cure rates in patients.

## Materials and Methods

### Experimental Design

The aim of this study was to build a scalable microfluidic culture chamber suitable for imaging twodimensional growth of bacterial microcolonies for long periods of time, and to develop and characterize a multi-condition microfluidic platform starting from this functional unit. We also aimed to validate the operation of the platform by implementing the first proof of concept of a single-cell dose-response assay with the tubercular pathogen under BSL3 containment.

### Bacterial strains and growth conditions

Plasmids cloning, sequencing and amplification steps were carried out in *Escherichia coli* TOP10 strains, grown at 37 °C under shaking conditions in LB medium containing the appropriate selective antibiotic: 100 μg/mL ampicillin; 50 μg/mL kanamycin; 100 μg/mL hygromycin. All mycobacterial strains were cultured at 37 °C under shaking conditions in Middlebrook 7H9 broth enriched with 0.5% BSA, 0.5% glycerol, 0.2% glucose, 0.085% NaCl, and 0.05% Tween-80. Middlebrook 7H10 agar plates were supplemented with 10% OADC and 0.5% glycerol. Mycobacterial transformants were selected on 20 μg/mL kanamycin, or a combination of 15 μg/mL kanamycin and 50 μg/mL hygromycin. For the construction of GyrA-mCherry fluorescent reporters, second recombination events were selected on Middlebrook 7H10 containing 5% sucrose for *M. smegmatis* or 2.5% sucrose for *M. tuberculosis*.

Bacterial stocks were prepared by adding 15% glycerol into late exponential-phase cultures (OD_600nm_ 1.0) obtained from single colonies, frozen at −80 °C, and used only once to start primary cultures (1:100 dilution). For final assays, primary cultures were grown up to mid-log phase (OD_600nm_ 0.5–0.8) and used to inoculate final secondary cultures (OD_600nm_ 0.025 dilution) until mid-log phase. For strains carrying chromosomal integrative plasmids, the selective antibiotic was maintained in both primary and secondary cultures, but removed during microscopy experiments.

### Strains construction

Translational fluorescent reporter strains were generated using a stable mCherry variant. An oligopeptide linker was used to fuse GyrA and mCherry, so as not to affect the folding of the native protein. Plasmids used for cloning were checked by sequencing and final constructs were confirmed by restriction enzyme profiling. Stable GFP variant under the control of a constitutive promoter was inserted into the L5 phage integration site (*13*). Mycobacteria were transformed by electroporation.

Fluorescent reporters of *M. smegmatis gyrA* (*MSMEG_0006*) and *M. tuberculosis gyrA* (*rv0006*) were obtained by two events of homologous recombination, using the suicide vector pJG1100 (*13*). In particular, the *mCherry* open reading frame was inserted into the *gyrA* chromosomal locus just upstream the stop codon of *gyrA* via an oligonucleotide sequence, generating C-terminus translational reporters. Amplicons of about 600 bp, upstream and downstream the chromosomal insertion site, were ligated to mCherry using compatible restriction sites, and cloned into pJG1100. After electroporation, the first recombination event was selected on antibiotics, while the second recombination event was selected on sucrose, and both events were confirmed by PCR analysis of genomic DNA (Figs. S5A and S6A). The final GyrA-mCherry reporters were named GMS5 for *M. smegmatis* and GMT18 for *M. tuberculosis*. GMT18 was further transformed with a plasmid constitutively expressing GFP, and the final double fluorescent reporter was named GMT18-GFP (Table S1).

### MIC assessment by resazurin assay

Secondary cultures were diluted to OD_600nm_ 0.005 using pre-warmed Middlebrook 7H9 medium without detergent, and 100 μL of bacterial suspension were dispensed into a 96-well plate, except the well with the highest concentration of drug to be tested that was filled with 200 μL of cell suspension. The drug was added to this well and then two-fold serially diluted. Plates were incubated at 37 °C for 24 h for *M. smegmatis* and 7 days for *M. tuberculosis*, then 10 μL of 0.01% resazurin were added to each well and incubated for 24 h at 37 °C, before visual quantification. The lowest drug concentration causing cell mortality, assessed as blue color, accounted for the MIC.

### Microfabrication

The fabrication of the microfluidic devices consisted of the following steps: mask design and printing; wafers fabrication; soft lithography; PDMS layers assembly and PDMS-glass bonding. Masks were designed using AutoCAD 2017, and low-resolution photomasks were printed at Selba S.A. (Data S1).

For each device, two micropatterned silicon wafers were created, one for the flow layer (FL-W) and the second one for the control layer (CL-W). Before coating, standard silicon wafers were dehydrated at 200 °C on a hot plate for 30 min. The FL-W was produced by spin-coating SU8-2025 photoresist onto a wafer at 3000 rpm for 30 sec, to generate a 30-μm thick layer, and soft-backed at 65 °C for 1 min, followed by incubation at 95 °C for 5.5 min. The photoresist was crosslinked under 155 mJ/cm^2^ UV light for 11.5 sec, and the wafer backed at 65 °C for 1 min, followed by incubation at 95 °C for 5 min. The wafer was developed by immersion into propylene glycol methyl ether acetate (PGMEA) for 4 minutes and washed with isopropanol. The CL-W was generated by spin-coating the SU8-2100 photoresist onto a standard silicon wafer at 1500 rpm for 30 sec, to generate a 200-μm thick layer, and soft-backed at 65 °C for 5 min, followed by incubation at 95 °C for 20 min. The photoresist was crosslinked under 240 mJ/cm^2^ UV light for 21.8 sec, and the wafer backed at 65 °C for 5 min, followed by incubation at 95 °C for 10 min. The wafer was developed by immersion into PGMEA for 15 minutes and washed with isopropanol. The FL-W and CL-W were hard-backed at 180 °C for 2 hours, and silanized with a drop of Trichloro (1H,1H,2H,2H-perfluorooctyl) silane under a vacuum chamber overnight. The height of the microchannels was confirmed using a profilometer DektakXT (Bruker).

The FL-W and CL-W were used as molds for PDMS soft lithography. To promote bonding between the two layers, we used a different amount of cross-linking agent, which diffuses from the layer with the highest concentration to the layer with the lowest concentration. The FL-W was covered with a degassed mixture of 20 g of silicone elastomer SYLGARD 184 (Dow Corning) and 1 g of cross-linking agent (20:1 ratio), spin-coated at 1500 rpm for 60 sec to obtain a 50-μm thick layer, incubated at room temperature for 20 min on a flat surface, and baked at 80 °C for 18 minutes. The CL-W was placed into a square Petri dish and covered with a degassed mixture of 40 g of silicone elastomer and 8 g of crosslinking agent (5:1 ratio), further degassed for 30 min in a vacuum chamber, baked at 60 °C for 30 minutes, and the patterned PDMS area was cut with a blade and gently separated from the mold.

The inlet ports were generated using a biopsy punch (1.0 mm OD, Harris Uni-Core). The PDMS CL was manually aligned with the FL, using the circular chambers in the two layers as references. Metallic connectors (0.8/1.2 mm ID/OD, Phymep) were inserted into the inlet port of the top element. Finally, the merged FL and CL were immersed in a degassed mixture of 30 g of silicone elastomer and 3 g of cross-linking agent (10:1 ratio), placed in a vacuum chamber for 30 min until degassing was completed, and baked at 80 °C overnight. The whole PDMS assembly was gently separated from the FL-W, the metallic connectors were unplugged from the inlet and outlet ports of the CL, and the inlet and outlet ports of the FL were generated with a biopsy punch (1.0 mm OD).

The PDMS assembly was bonded with a glass coverslip by oxygen plasma treatment. For the single-condition device we used a 24 x 50 x 0.175 mm coverslip, and for the 5-condition platform we used a 40 x 60 x 0.175 mm coverslip. Coverslips were cleaned with isopropanol prior to plasma treatment. Both the coverslip and the PDMS assembly, with the microchannels facing upward, were placed inside the plasma chamber. Initially the vacuum was created until 0.1 mbar, then oxygen was injected until a pressure of 0.25 mbar was reached, and the treatment was maintained for 30 sec for the single-condition device, and for 60 sec for the 5-condition device. Upon plasma activation, the PDMS assembly was placed on the coverslip and incubated at 80 °C overnight to complete the bonding. For decontamination, the device was exposed to ozone/UV treatment for 15 min inside a UVO-Cleaner (Jelight) and stored in a sterile 20-mm petri dish.

### Characterization of PDMS membrane deformation

The deformation of the PDMS membrane was evaluated in the single-condition device, using a 100 μM solution of fluorescein (FITC) perfused into the FL at 50 mbar, and applying a pressure range from 55 to 80 mbar in the CL with incremental steps of 1 mbar. The FL inlet was connected to the FITC reservoir and the FL outlet to a waste bottle. The CL inlet was connected to a water reservoir, and the CL outlet to a short tubing, which was closed with a plier upon water injection. The pressures in the CL and FL were fine-tuned via a flow controller (Fluigent). FITC fluorescence (Ex 475/28, Em 525/48 nm), 10% T and 10 msec exposure time was acquired from each microchamber at 30 sec intervals, using an inverted DeltaVision Elite Microscope (Leica) equipped with an APOCHROMAT 20x/1.0 air objective (ZEISS) and a high-speed sCMOS camera, 2560 x 2160 pixels, pixel size 6.5 x 6.5 μm, 15-bit, spectral range of 370–1100 nm. Image stacks were analysed on ImageJ 1.53a, tracing a line along the diameter of each microchamber and acquiring the fluorescence intensity over time with the Plot Profile function. The background mean fluorescence intensity was extracted from areas of the device without liquid circulation and subtracted to the microchamber measurements. Fluorescence decrease in the microchambers was considered to be proportional to increasing pressure in the CL.

### FITC gradients generation and quantification

Three 125 mL bottles were prepared, two containing 7H9 medium and the third containing a 100 μM FITC solution in 7H9 medium, and were connected to a bidirectional M-switch valve (Fluigent) with 10 inlet and 1 outlet port. One medium and the FITC bottle were connected to two separate inlet ports of the valve, and the outlet port of the valve was connected to the left inlet of the device. The right inlet port of the device was connected to the third bottle of medium. The outlet port of the device was connected to a flowmeter (Fluigent) and then to an empty waste bottle. The switch between conditions was achieved via the ESS Control software (Fluigent). A Tygon tubing was used to connect the CL inlet to a water bottle, and the CL outlet was connected to a short tubing. The three bottles were driven by the first channel (C1) of the flow controller (Fig. 2A). The M-switch was set on medium perfusion mode. The water bottle was driven by the second channel (C2) and the waste by the third channel (C3). The pressure was set at 100 mbar on C1 and C2 and, once the CL and FL were filled with liquids, their outlet tubing was closed with pliers and the pressure was increased to 400 mbar on C1. After 20 min of incubation to remove bubbles, the plier closing the FL outlet was removed. The pressure was set at 100 mbar in C1, 30 mbar in C2, and 20 mbar in C3. Based on the measured flow rate in the FL, the pressure was fine-tuned to 150 μL/h in C1. To verify the formation of different gradient patterns, FITC fluorescence was acquired from each microchamber over time.

To generate static gradients (Fig. 3B), 7H9 medium and FITC solution were injected alternately for targeted periods of time by switching the position on the M-switch valve, in the absence of capacitors. To generate pulsing (PK-like) gradients (Fig. 3C), two capacitors were integrated just upstream the inlet ports of the FL. Capacitors were 1-cm Tygon tubing (2 mm ID, BioRad), having Luer connectors on each extremity. The pressure in the FL was fine-tuned to 120 μL/h on 7H9 medium mode. To initiate a single pulse, the M-switch valve was moved to the FITC position for 12 min and switched back to 7H9 medium position for 12h. To generate sequential pulses, the M-switch valve was moved to FITC position for 12 min, switched back to 7H9 medium position for 6h, and then the sequence was repeated. Fluorescence imaging was carried out as described above, and pixel intensity was measured on regions of interest (ROI) of equal size and subtracting the background intensity from areas of the device without liquid circulation.

### Device assembly, cell loading and fluids management

To make the PDMS-glass devices suitable for cell loading and tracking by time lapse microscopy, different tubing, interfaces and fittings were prepared and autoclaved prior to final assembly (Fig. S2).

Inlet ports of the FL were used for medium and drug perfusion from the reservoir bottles, which were closed with pressure lids (Fluigent), and the flangeless fittings (IDEX Health & Science) of these lids were connected to 50-cm Teflon tubing. In turn, the inlet ports were connected to 5-cm Tygon tubing (Saint-Gobain), connected to a metallic fitting at the other end. Then Teflon (TFE) and Tygon tubing were joined with a piece of silastic and sealed with PDMS, in the absence or presence of a capacitor. The outlet port of the FL, used both to seed the cell suspension and to drain waste fluid into the waste bottle, was connected to a tripartite tubing as follows. A PVDF Y-connector (Buerkle) was connected to two 10-cm Tygon tubing and one 50-cm Tygon tubing (1.2 mm ID, Saint-Gobain). The Tygon tubing used for cell loading was connected to a female Luer-to-barb connector and to a Luer lock (Cole Parmer). The other Tygon tubing was plugged into the device via a metallic fitting. The longer tubing was connected to a 250 mL waste bottle closed with a pressure lid (Fluigent) via flangeless fittings. A flowmeter (Fluigent) was also integrated between the outlet tubing and the waste bottle, to maintain the flow rate constant.

The inlet port of the CL was connected to a 50-cm Tygon tubing, having a metallic fitting at one end and a flangeless fitting at the other end, plugged into the water reservoir. The outlet port of the CL was connected to a 5-cm Tygon via metallic fitting and closed with a plier.

To load bacterial cells, the outlet tubing was closed with a plier. Exponentially growing cells were disaggregated through a pre-equilibrated 5-μm filter (Merck), and about 400 μL of cell suspension were gently injected into the *ad hoc* Tygon tubing by hand, using a 1-mL syringe. When loading pathogenic *M. tuberculosis* cells, 0.2 μm filters were also added at the extremities of the inlet tubing, to prevent the pathogen from coming out of the device during the loading procedure. Next, the device was incubated at room temperature for 10 min. The entire procedure was performed under a class II biosafety cabinet, and the waste bottle was filled up for one fifth with a 2.6% sodium hypochlorite solution. At the end of the incubation period the syringe was unplugged, replaced by the Luer lock, and the junction was sprayed with mycobactericidal disinfectant. Finally, the device was connected to the reservoir and waste bottles and mounted on the microscope stage (Figs. 2A and S2).

To generate the drug gradient, one sterile reservoir bottle was filled with 7H9 medium, the second one with 7H9 medium supplemented with 250 ng/mL of moxifloxacin (5X-MIC). These bottles were connected to the first and second port of an M-switch valve, in turn connected to the inlet port ‘c’ of the device. The third reservoir bottle was filled with 7H9 medium, and connected to the inlet port ‘d’ of the device (Fig. 2A). Fluids were controlled from the multi-channel ESS Control software via pressure caps: C1 drove the device inlet of the FL, C2 drove the CL, and C3 drove the device outlet of the FL. After cell seeding, the CL was filled with water at 100 mbar and closed with a plier. The FL was perfused with 7H9 medium at 200 mbar for 20 min and the outlet closed with a plier for other 20 min to degas the pipe. Next, in the single condition device, C1 was set at 60 mbar and C2 was set at 30 mbar, targeting a final flowrate of about 100 μL/h. In the 5-condition platform, C1 was set at 60 mbar and both C2 and C3 were set at 30 mbar, targeting a final flowrate of about 150 μL/h. To inject the drug, the M-switch valve was moved to the second position, and the pressure was set at 60 mbar on C1, 30 mbar on C2, and 20 mbar on C3. At the end of drug injection, the M-switch valve was moved to the initial position.

### Time-lapse imaging

The assembled device was mounted on the stage of our inverted DeltaVision Elite Microscope, equipped with a UPLFLN100XO2/PH3/1.30 oil objective (Olympus), using a custom-made metal support (Fig. S3). Exposure conditions for *M. smegmatis:* phase contrast 100% T, 150 msec; mCherry (Ex 575/25, Em 625/45 nm) 50% T, 150 msec. Exposure conditions for *M. tuberculosis*: phase contrast 100% T, 150 msec; FITC (Ex 475/28, Em 525/48 nm) 32% T, 10 msec; mCherry (Ex 575/25, Em 625/45 nm) 50% T, 250 msec. In each microchamber between 5 and 30 x,y,z coordinates were recorded, each one containing from 3 to 10 bacilli. The image acquisition rate was set at intervals of 20 or 30 min for *M. smegmatis* and of 3 h for *M. tuberculosis*. At the end of the experiment, the entire platform was washed with 2.6% sodium hypochlorite, the device was separated from tubing and discarded into a sharp waste container. The electronic M-switch valve and the flowmeter were extensively washed with 2.6% sodium hypochlorite, 70% ethanol and sterile water for 1 hour. Tubing, fluidic interfaces and fittings were placed into plastic bags, sprayed with mycobactericidal disinfectant and autoclaved. Each experiment was performed at least twice independently.

### Microcolony analysis

The growth of individual microcolonies from the single condition device was assessed by manual segmentation over time. The analysis of microcolonies from the 5-condition device was carried out in a semi-automated manner (Fig. S5B and Data S2-S7). Image stacks were separated by channel using a customized macro (Ch-Extraction.ijm). To generate binary masks, a specific fluorescence channel was used: mCherry for *M. smegmatis* (mCherry-Mask.ijm), and FITC for *M. tuberculosis* (GFP-Mask.ijm). Images were first smoothed and then the Auto threshold plugin was used to distinguish the foreground from the background pixels. The Huang method (*79*) was used for the mCherry channel, and the Intermodes method (*80*) was used for the FITC channel. The binary masks were used to measure area and mean fluorescence from the ROI (ROI-Measure.ijm), and the inverse of the binary masks were used to measure the background fluorescence (BG-Measure.ijm), to be subtracted from the corresponding ROI. Data were saved as .csv files, which were merged and processed, using a customized R script (csv-Management.R). The microcolony coordinates were also recorded from the image metadata files, in order to determine the position relative to the center of the microchamber.

### Single-cell analysis

*M. tuberculosis* single-cell area and mean fluorescence were measured from independent ROI using the Selection Brush Tool and the ROI Manager macro at given time points. In each image, the background mean fluorescence was also measured and subtracted from the fluorescence of individual cells, which were segmented in the corresponding image.

*M. tuberculosis* single-cell counting over time was performed using the Cell Counter plugin, and scoring the total cell count 24 h before drug exposure; dividing cells (alive); lysed cells (dead); GFP-dim cells (likely dead); cells that resumed growth after drug exposure (alive); and non-growing GFP-positive cells (likely viable). Quantified subpopulations were expressed as a function of time and as fractions.

### Calculation of parameters

The growth rate of individual microcolonies was measured by nonlinear regression analysis of the area increase as function of time, fitting an exponential growth equation: Y=Y0*exp(*k*\*t), where Y0 is the initial microcolony size (μm^2^) and *k* is the rate constant (time^-1^). The doubling time is ln2/*k*.

The AUC and associated parameters were measured based on the highest concentration of moxifloxacin injected, according to the dilution factors assessed from the formation of FITC gradients, and using the MIC concentration (50 ng/mL) as the baseline.

Dividing and lysed cells were counted starting from the number of cells 24 h before drug exposure, and progressively adding the number of divisions and subtracting the number of lysis events, obtaining the total count throughout the experiment. Next, the number of division events was divided by the cell count at each time point and by the image acquisition time frequency, obtaining the division rate per hour.

The relative IC50 was computed from the fraction of viable cells at different moxifloxacin doses, by interpolating a four-parameter dose-response curve according to the variable-slope model. We considered viable those cells that were non-growing but GFP positive and those that resumed growth.

### Statistical Analysis

Plots and statistical analyses were generated using Prism 9 (GraphPad). One-way ANOVA and multiple comparisons tests were used to assess significant differences within datasets containing more than two groups. Linear relationship between *x,y* datasets was determined by computing the Pearson correlation coefficient. Plots include datasets derived from at least two independent experimental replicates. *N* values, *P* values and specific statistical tests are indicated in each legend.

## Acknowledgments

We thank Pietro Slavich (LPTHE) for reading of the manuscript and critical discussions. We thank Théo Guillerm for his contribution in analyzing the gradient formation with dyes. We thank Gaël Millot (Bioinformatics and Biostatistics Hub of the Institut Pasteur), for support in writing the R script for managing .csv files. We thank Samy Gobaa for providing us access to the Biomaterials & Microfluidics core facility of the Institut Pasteur.

## Author contributions

Conceptualization: GM, MM; Methodology: MM, GM; Investigation: MM, NG, GM; Resources: MM, GM; Data curation: MM; Formal Analysis: MM, NG, GM; Visualization: MM, GM; Writing—original draft: GM; Writing—review & editing: MM, NG, GM; Supervision: GM; Project Administration: GM; Funding Acquisition: GM.

## Data and materials availability

All data needed to evaluate the conclusions in the paper are present in the paper and/or the Supplementary Materials. Additional data related to this paper may be requested from the authors.

## Supplementary Figures

**Fig. S1.**
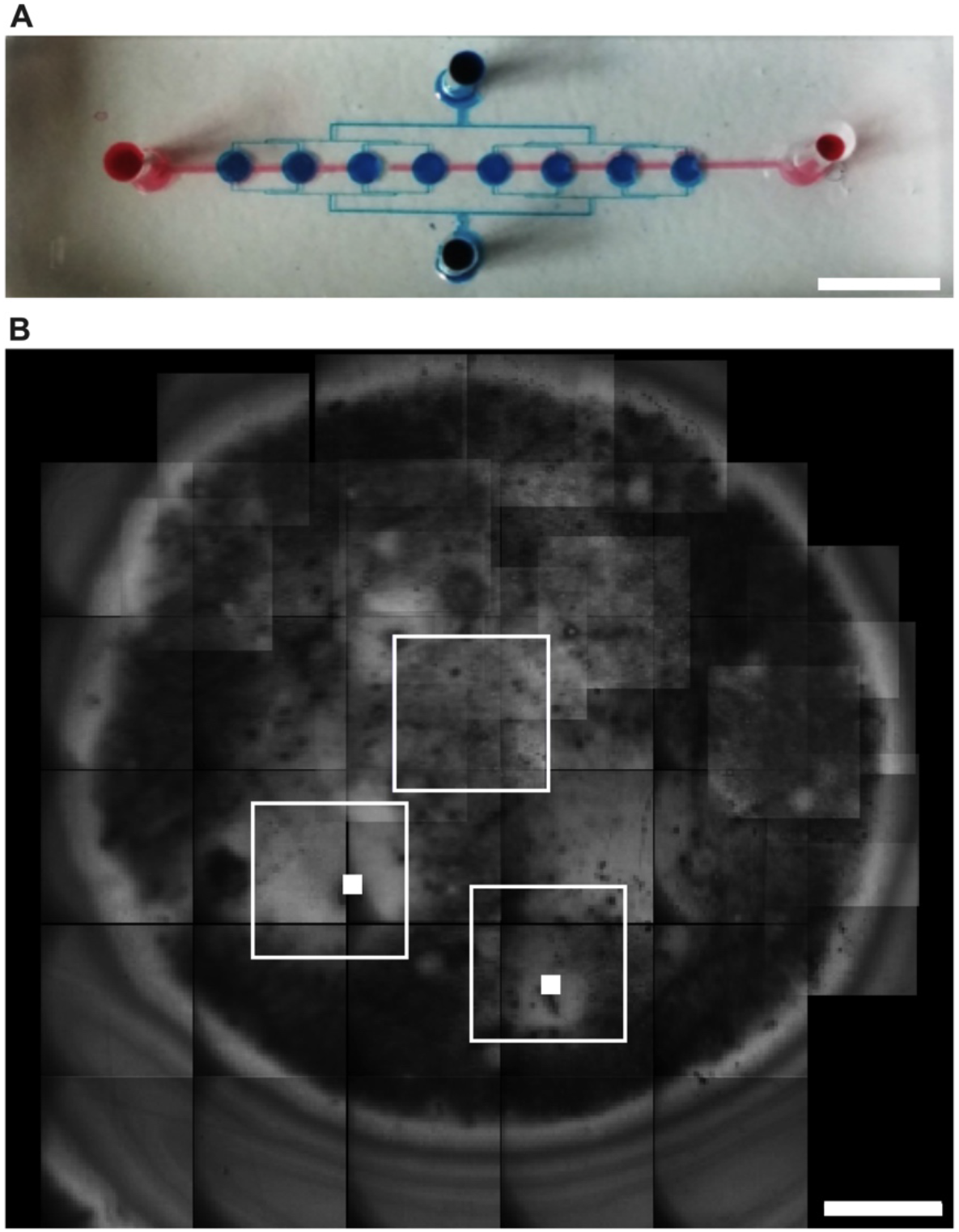
Single-condition prototype. (**A**) Picture of a representative single-condition device, where eight microfluidic culture chambers are connected by a tree-shaped flow layer (FL, blue dye) and overlaid by the control layer (CL, red dye). Scale bar, 3.5 mm. (**B**) Representative mosaic picture of the 2D-growth area formed between the PDMS membrane and the coverslip by applying a pressure of 25 mbar in the CL from the pressure controller. The field of view is excited at 475/28 nm and images are acquired in bright field. Scale bar, 50 μm. The brightest areas indicate the presence of a liquid interface (empty squares). *M. smegmatis* microcolonies are also marked (filled squares).

**Fig. S2.**
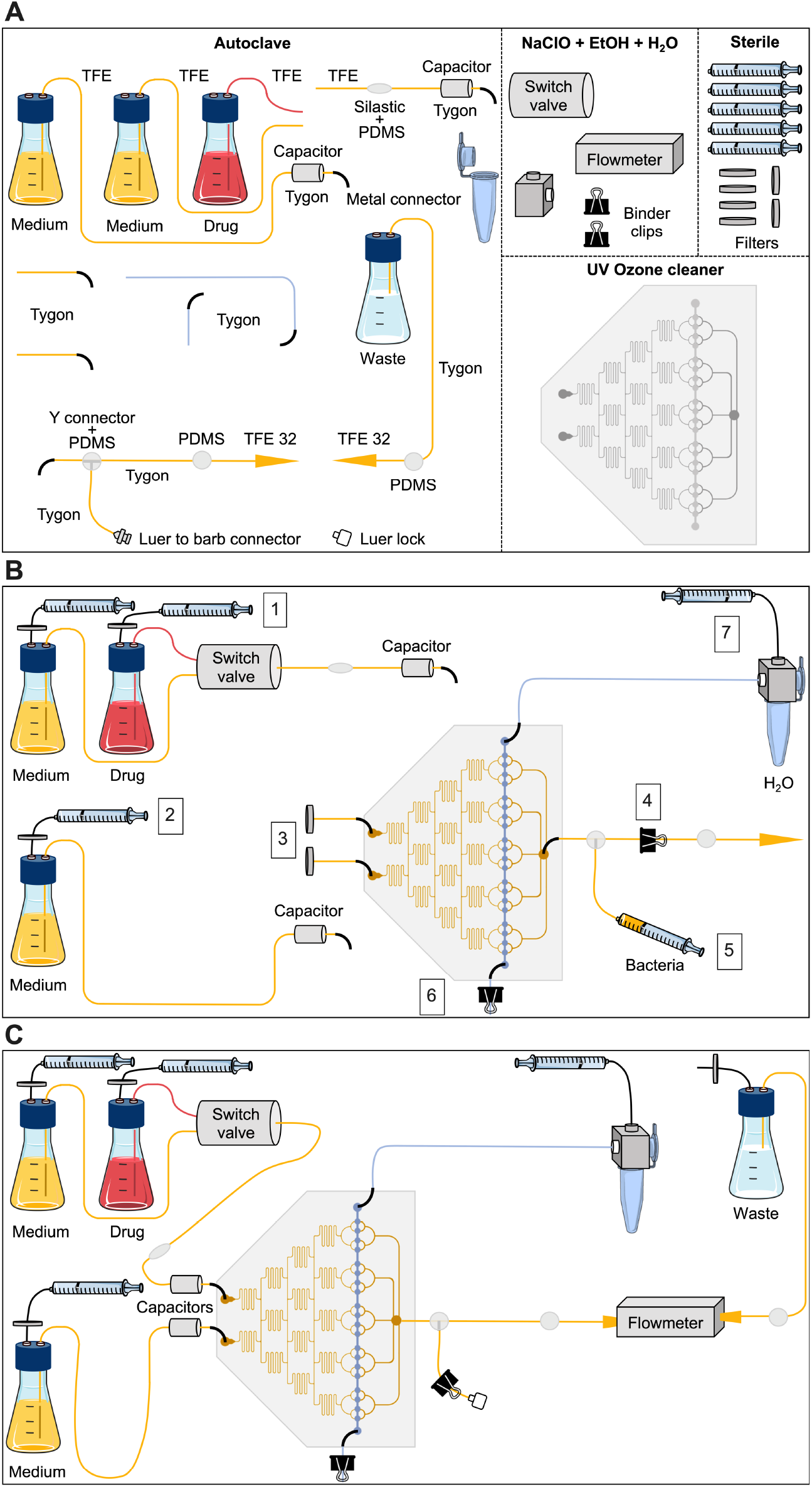
Preparation and assembly of the five-condition platform. (**A**) Decontamination of the different components of the five-condition platform. The color of the tubing distinguishes the application: culture medium in the FL (yellow); culture medium containing the highest drug concentration in the FL (red); water in the CL (blue). Materials constituting the different components are indicated. Cured PDMS is used to seal critical connections between tubing. Dashed lines separate different decontamination methods. Except for bottles, tubing and connectors, all other components are not autoclavable. Electronic systems (Fluigent) are perfused with 0.5% bleach, followed by 70% alcohol and sterile deionized water. The microfluidic device is decontaminated by UV and ozone treatment, and disposed after use. (**B**) Priming of the different components of the five-condition platform is carried out under a class II biosafety cabinet. To prevent air accumulation in the fluidics, medium and drug bottle reservoirs and tubing are pre-filled by manually pushing air from 10 mL syringes into bottles closed with a cap having two ports. One medium bottle and the drug bottle are connected to the M-switch valve (Fluigent), in turn connected to the tubing with the capacitor. The second medium bottle is connected to a separate tubing with the second capacitor. Capacitors are 1 cm long Tygon tubing with an inner diameter of 2 mm and are connected via Luer fittings on each extremity. The whole tubing network is filled with medium. For loading bacteria, the inlet ports of the device are plugged with Tygon tubing through two metal connectors. The bacterial suspension contained in a syringe is loaded through the outlet port via a secondary tubing and leaving the primary outlet tubing clamped with a binder clip. Next, water is manually perfused into the inlet of the CL, after closing the CL outlet with a binder clip. Numbers indicate consecutive actions. (**C**) Final assembly of the five-condition platform. After 10 min of incubation, the syringe containing the bacterial suspension is removed, and the secondary outlet tubing closed with a binder clip, disinfected, closed with a Luer lock and disinfected again. The bottle reservoirs are connected to the inlet ports of the device, and the outlet tubing is connected to the flowmeter (Fluigent) and to the sealed waste, after removing the binder clip. The system assembly is ready to be connected to the flow controller and to be mounted on the microscope stage.

**Fig. S3.**
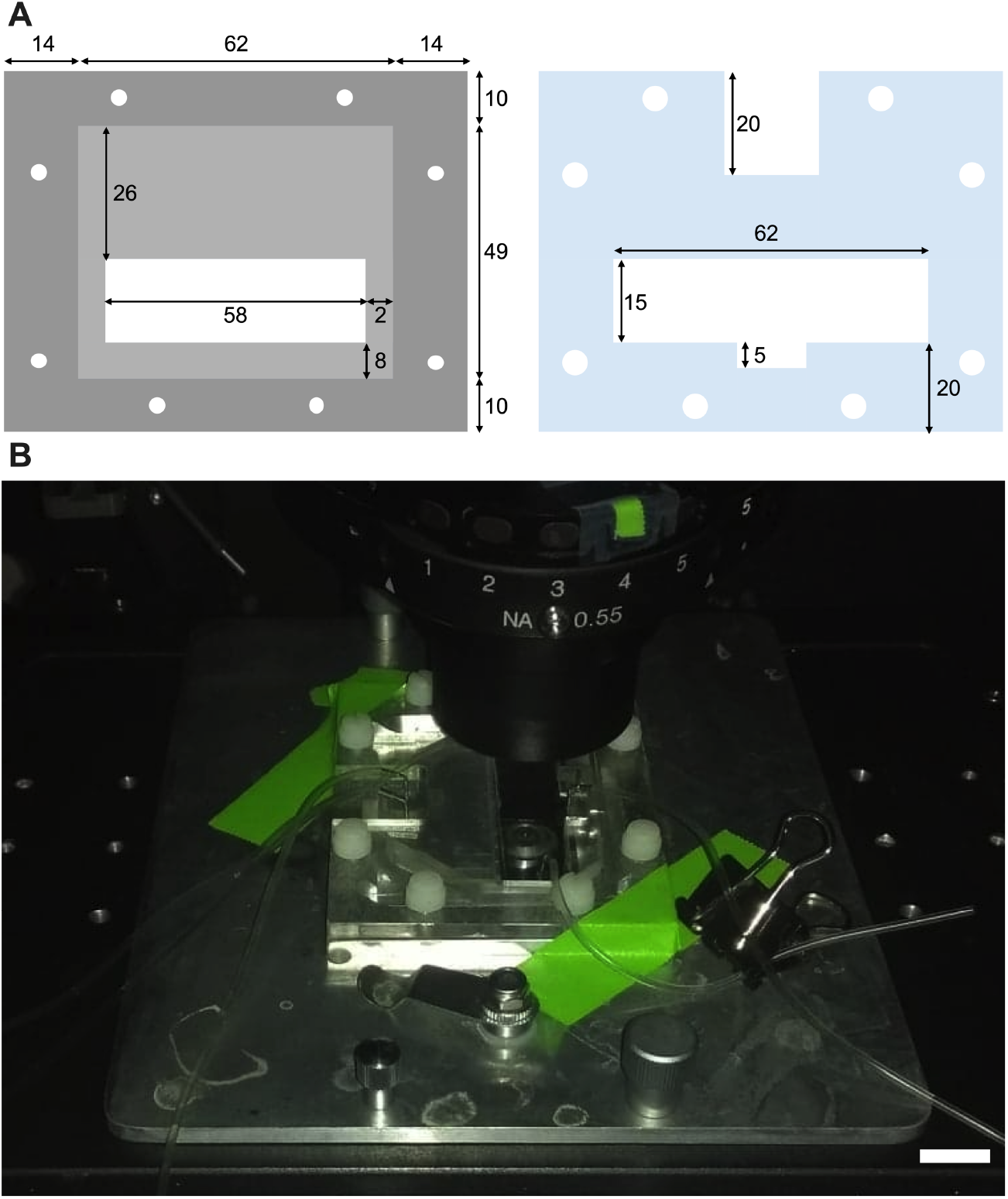
Mounting of the 5-condition device. (**A**) Schematic of the metal and acrylic holders used to mount the five-condition device on the microscope stage. Circular holes are used to secure the system with plastic screws. Numbers indicate dimensions in millimeters. (**B**). Picture of the microscope stage with the five-condition device fixed with metal clips and tape. Scale bar, 10 mm.

**Fig. S4.**
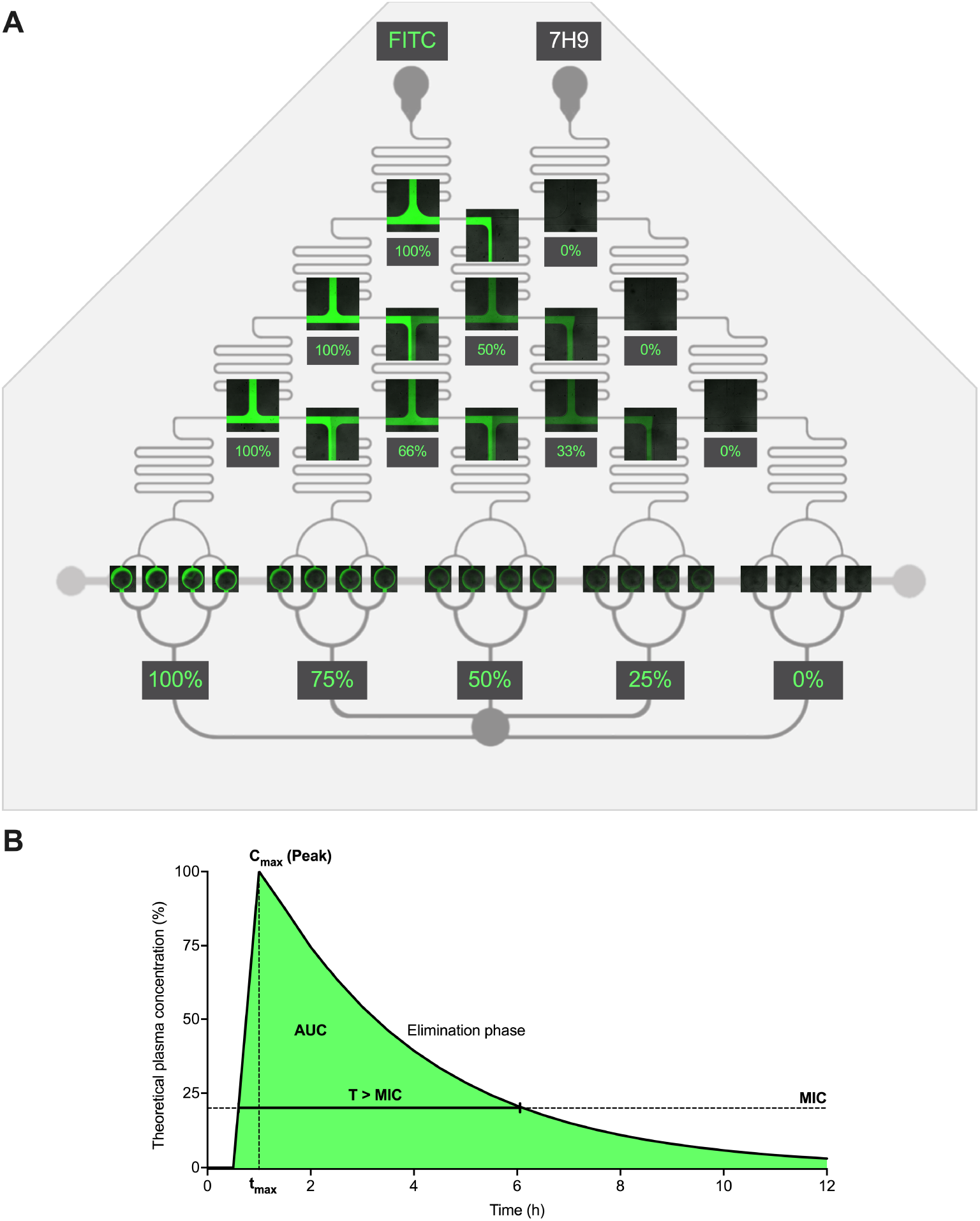
Generation of a static linear concentration gradient and PK parameters. (**A**) Representative images of serpentine junctions and exits and of the microchambers, overlaid on the plan of the 5-condition device, during steady-state injection of a FITC solution (100 μM) into the left inlet-port and of 7H9 into the right inlet-port, at a flow rate of 120 μL/h. The dilution rates shown in the figure are the theoretical ones, derived from the assumption of a linear gradient. Bright field and fluorescence images are merged. Microchannels thickness, 0.3 mm, and microchambers diameter, 1 mm. (**B**) Schematic of a theoretical drug PK profile, indicating MIC; peak plasma concentration (C_max_); time to reach the C_max_ (t_max_); area under the curve (AUC); and time over the MIC (T > MIC).

**Fig. S5.**
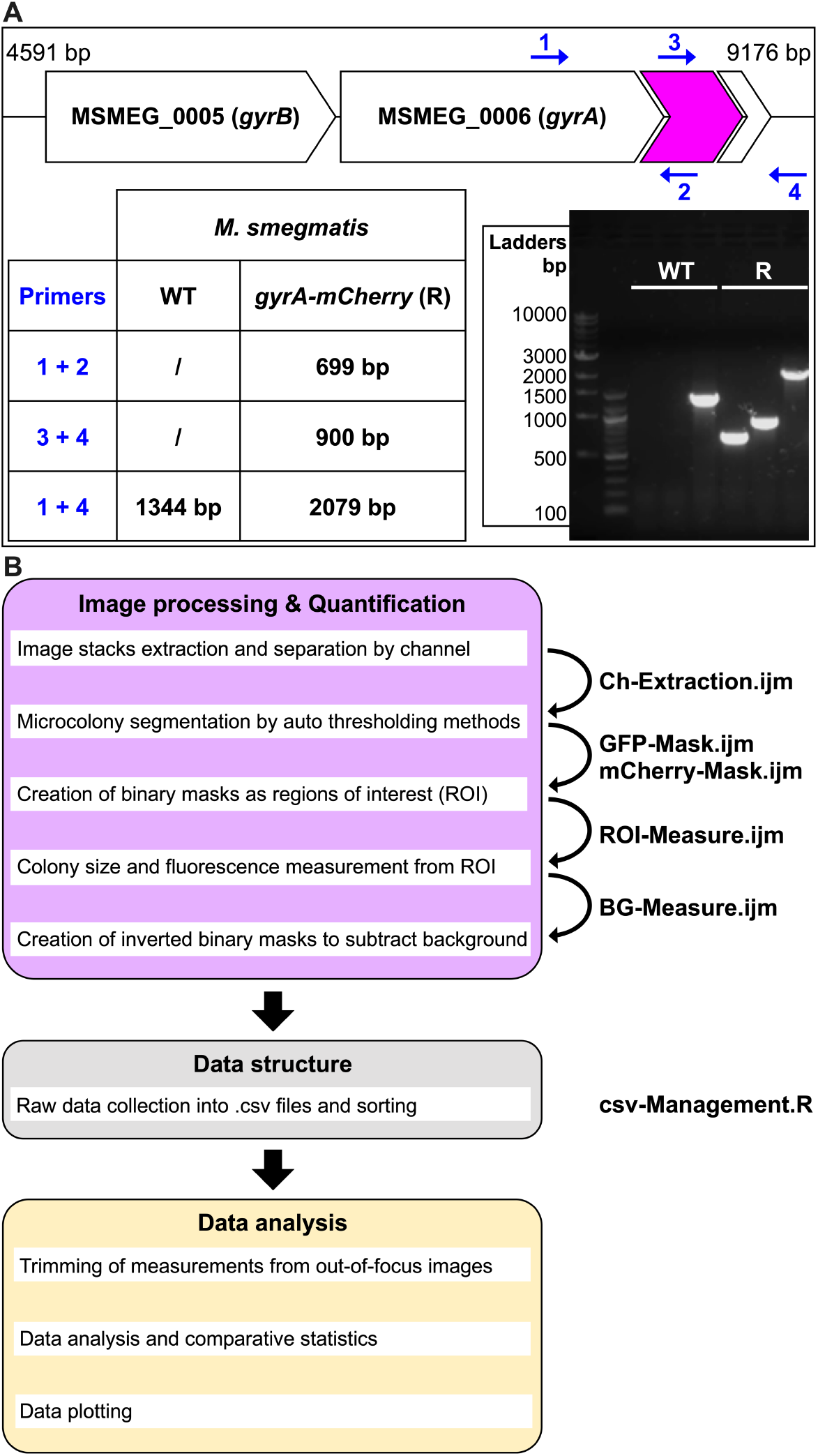
GyrA-mCherry reporter construction and image analysis workflow. (**A**) Schematic of DNA gyrase encoding operon in *M. smegmatis* mc^2^155 genome, with *mCherry* gene (magenta arrow) inserted in frame, just upstream of *gyrA* stop codon. Blue arrows represent the primers used for PCR analysis, to check *mCherry* chromosomal insertion. Expected amplicon sizes are indicated in the table for wild type (WT) and reporter (R) strains, and shown in a 1% agarose gel. (**B**) Data analysis diagram separated into three steps: i. Extraction of image stacks from time-lapse movies, creation of masks and microcolony data extraction from ROI and inverted ROI (five ImageJ macros, lavender box); ii. Extraction of .csv files containing microcolony-derived parameters over time (area, mean fluorescence and standard deviation of fluorescence) and corresponding background measurement for each image (one R script, gray box); iii. Removal of values from out-of-focus images, data processing, statistical analysis and data visualization (Excel and GraphPad Prism, yellow box).

**Fig. S6.**
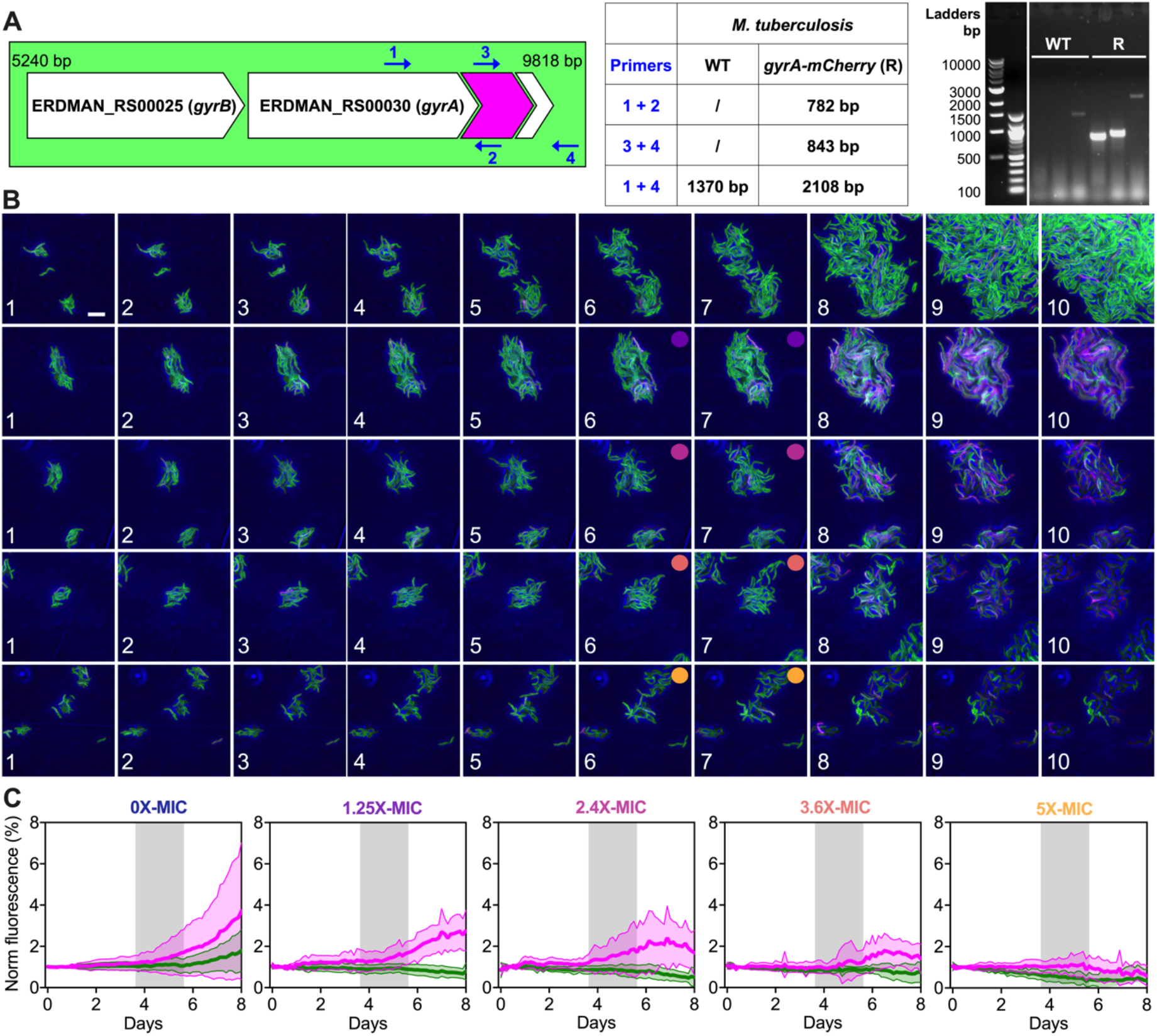
Multi-condition and multi-phasic time-lapse microscopy of *M. tuberculosis* GFP_cyt__GyrA-mCherry reporter. (**A**) Schematic of DNA gyrase encoding operon in *M. tuberculosis* Erdman genome, with *mCherry* gene (magenta arrow) inserted in frame, just upstream of *gyrA* stop codon. Green color indicates constitutive expression of a green fluorescent marker in the cytosol (GFP_cyt_). Blue arrows represent the primers used for PCR analysis, to check *mCherry* chromosomal insertion. Expected amplicon sizes are indicated in the table for wild type (WT) and reporter (R) strains, and shown in a 1% agarose gel. (**B**) Time-lapse image series of exponentially growing *M. tuberculosis* seeded into the five-condition device and stressed with a stable gradient of moxifloxacin. Each row is representative of a chamber group: absence of drug is unlabeled and colored circles indicate different moxifloxacin concentrations (purple: 62.5 ng/mL; pink: 120 ng/mL; red: 180 ng/mL; yellow: 250 ng/mL), according to the dilution factors measured in Fig. 3B. Phase-contrast and fluorescence channels are merged. Images were acquired every 3 hours and numbers represent days. Scale bar, 5 μm. (**C**) Normalized microcolony GFP_cyt_ (green) and GyrA-mCherry (magenta) fluorescence during time-lapse microscopy. Datasets are expressed as mean ± SD (13 ≤ *N* ≤ 45 microcolonies, from two independent experiments). Concentrations (gray shadings) are relative to the MIC (50 ng/mL).

**Table S1.**
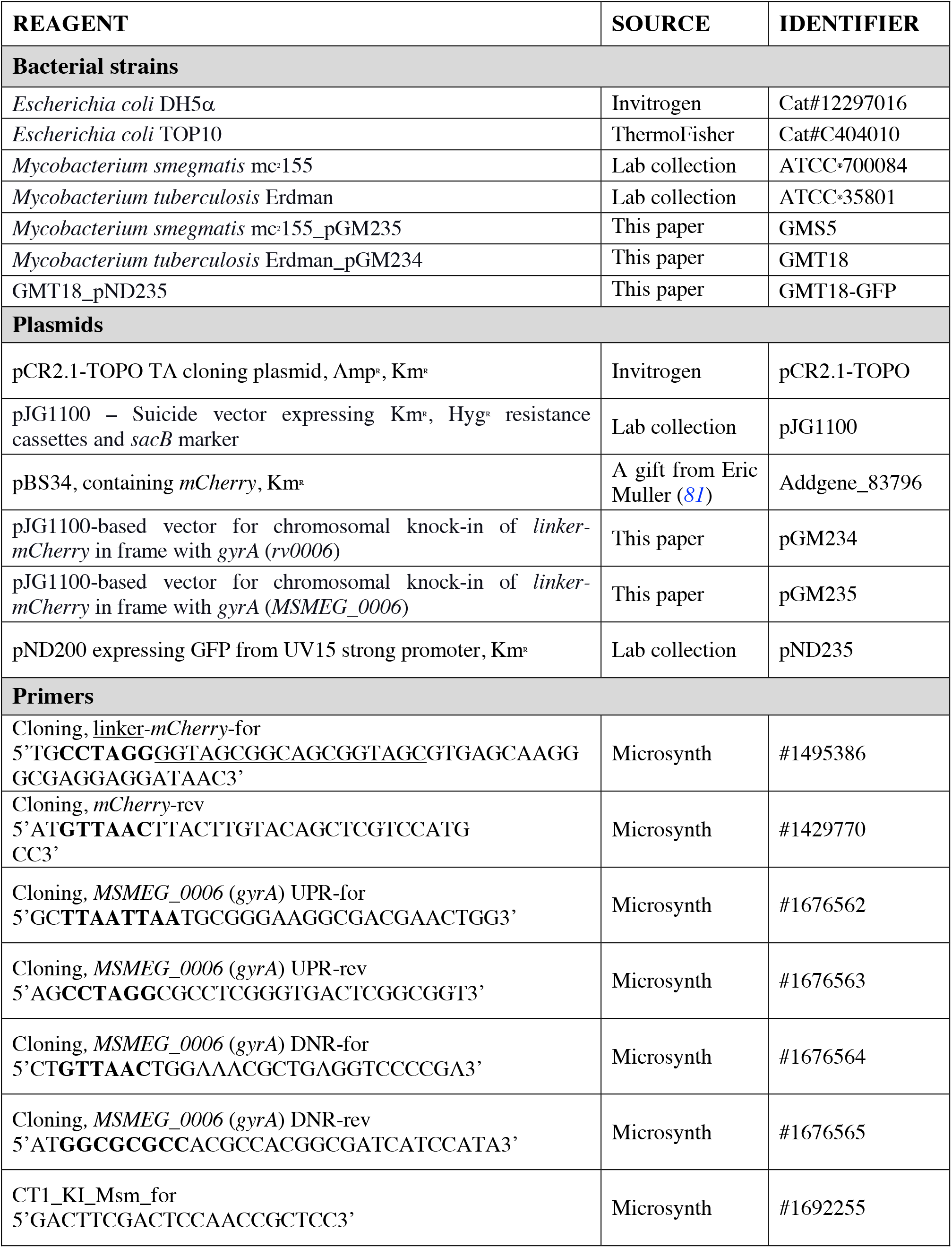
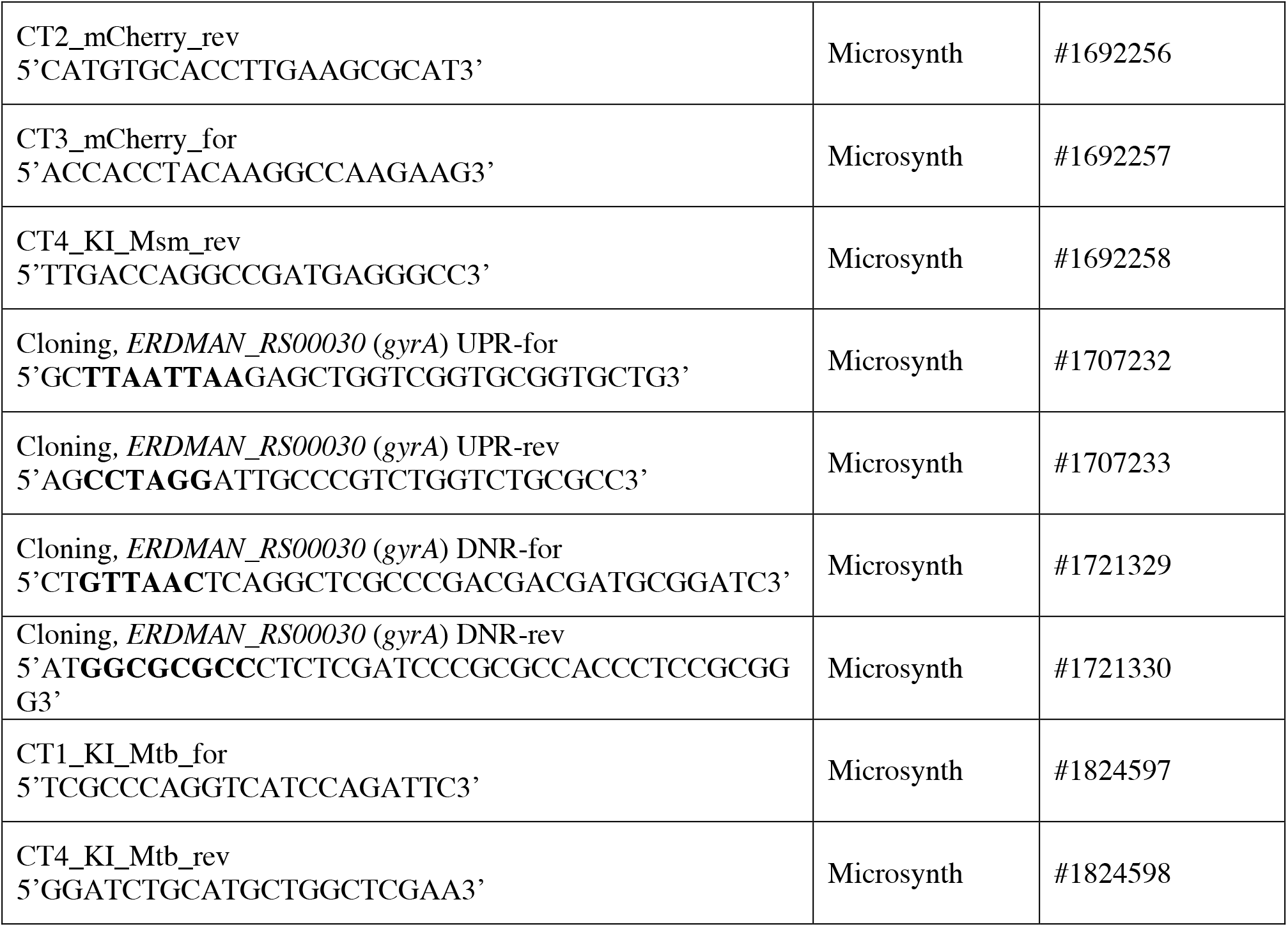
Strains, plasmids and primers used in this study. Resistance markers are indicated and restriction sites are bolded.

## Movies

**Movie S1. PDMS membrane actuation.** Representative movie of a microfluidic chamber, perfused with a FITC solution (100 μM), obtained with the ImageJ plugin Stack 3D Surface Plot. Fluorescence inside and outside the microchamber was acquired at constant flow in the FL (150 μL/h) and at incremental pressure steps in the CL, and was represented by a heatmap as a function of the microchamber size. Decreasing fluorescence represents lowering of the PDMS membrane.

**Movie S2. Combined time-lapse microscopy of *M. smegmatis* GyrA-mCherry reporter treated with a pulsing gradient of moxifloxacin.** Representative movies of exponentially growing bacteria seeded into different microchambers of the 5-condition platform. Bacteria were first grown in fresh 7H9 medium for 6 hours, then one microchamber (left side) was left without drug and the other four microchambers were perfused with different pulsing concentrations of moxifloxacin (Moxi) for 12 hours. The peak concentrations are expressed relative to the MIC (50 ng/mL). Finally, fresh 7H9 medium was perfused everywhere for 6 hours. Images were recorded every 20 minutes (10 fps). GyrA-mCherry (magenta) and phase contrast (blue) channels are merged. Time in minutes and drug concentrations are indicated. Scale bar, 10 μm.

**Movie S3. Combined time-lapse microscopy of *M. tuberculosis* GFP_cyt__GyrA-mCherry reporter treated with a static gradient of moxifloxacin.** Representative movies of exponentially growing bacilli seeded into different microchambers of the 5-condition platform. Bacilli were first grown in fresh 7H9 medium for 5 days, then one microchamber (left side) was left without drug and the other four microchambers were perfused with different concentrations of moxifloxacin (Moxi) relative to the MIC (50 ng/mL) for 2 days. Finally, fresh 7H9 medium was perfused everywhere for 3 days. Images were recorded every 3 hours (10 fps). Constitutive GFP_cyt_ (green), GyrA-mCherry (magenta) fluorescence and phase contrast (blue) channels are merged. Time in hours and drug concentrations are indicated. Scale bar, 5 μm.

## Supplementary Data

**Data S1. AutoCAD mask design of 5-condition platform.** FL (up) and CL (down) are shown.

**Data S2. Ch-Extraction.ijm.** ImageJ macro for channels extraction from image stacks.

**Data S3. GFP-Mask.ijm.** ImageJ macro to generate binary masks from green fluorescence.

**Data S4. mCherry-Mask.ijm.** ImageJ macro to generate binary masks from red fluorescence.

**Data S5. ROI-Measure.ijm.** ImageJ macro to quantify different parameters from segmented images.

**Data S6. BG-Measure.ijm.** ImageJ macro to quantify the background fluorescence.

**Data S7. csv-Management.R.** R Studio script to sort datasets for analysis.

