## Supplementary material for "Dynamics of drug response in single mycobacterial cells by microfluidic dose-response assay": Data S1-S7: BG-Measure.ijm

```
//--------------------------------------------------------------------------------------------------------------------
// DATA SOURCES
//--------------------------------------------------------------------------------------------------------------------


ExpFile = "/Users/mmistret/Documents/Data/Microfluidic/Longterme_Imaging/05_Gradiant_Chip/03_5C_MTB/02_MOX/20210422_5C_MTB_250_MOX_R2/";

//OPEN ROI_MANAGER BEFORE TO START


//---------------------------------------------------------------------------------------------------------------------
// INPUT and OUTPUT folders
//---------------------------------------------------------------------------------------------------------------------

//ImageFluo to annalyse
GFP_file=  ExpFile + "GFP_TIFF/";

//ImageFluo to annalyse
MCherry_file=  ExpFile + "DSRed_TIFF/";

//Mask_Final
MaskVF_file =  ExpFile + "Mask_Merged/";

//Results GFP
Results_file_GFP =  ExpFile + "Results_GFP_bk_Merged/";

//Results MCherry
Results_file_MCherry =  ExpFile + "Results_mCherry_bk_Merged/";

//ROI
ROI_file= ExpFile + "ROI_bk_Merged/";

//--------------------------------------------------------------------------------------------------------------------
//--------------------------------------------------------------------------------------------------------------------


setBatchMode(true); 
GFP_list = getFileList(GFP_file);
//MCherry_list = getFileList(MCherry_file);
MaskVF_list = getFileList(MaskVF_file);
X=1;

for (i = 0; i <GFP_list.length; i++) {
        DataExtraction(GFP_file, GFP_list[i], X, MaskVF_list, MaskVF_list [i], Results_file_GFP, ROI_file);
       // DataExtraction(MCherry_file, MCherry_list[i], X, MaskVF_list, MaskVF_list [i], Results_file_MCherry, ROI_file);
        X = X+1 ; }
setBatchMode(false)


//--------------------------------------------------------------------------------------------------------------------
//--------------------------------------------------------------------------------------------------------------------

function DataExtraction(input, filename, X, MaskVF_list, Mask_filename, Results_file, ROI_file) {
		open(input + filename);	
        getDimensions(width, height, channels, slices, frames);
        open(MaskVF_file + Mask_filename);
       	selectWindow(Mask_filename);
        getDimensions(width, height, channels, slices, frames1);
		print (frames);
		print (filename);
		print (Mask_filename);
		run("Set Measurements...", "area mean standard display redirect=None decimal=1");

		for (j = 1; j <=frames ; j++) {
    		  selectWindow(Mask_filename);
  			  run("Measure");
  			  getRawStatistics(area, mean);
  			  //print (mean);
  			 if (mean == 255) {
  			 	selectWindow(Mask_filename);
  			 	run("Invert","slice");
  			 	}
  			 selectWindow(Mask_filename);
  			 run("Create Selection");
			 run("Measure");
			 getRawStatistics(area);
			 print(area);
			 if (area > 589820) { //depanding of IMAGE
				//run("ROI Manager...");
				//roiManager("add");	
			    selectWindow(Mask_filename);
			 	run("Next Slice [>]");	 
				selectWindow(filename);
				run("Next Slice [>]");	
			  	}
			 else {
				 //run("Make Inverse");
				 run("ROI Manager...");
				 roiManager("add");
				 selectWindow(Mask_filename);
				 run("Next Slice [>]");
			     selectWindow(filename);
				 run("Next Slice [>]");
				 //print (j);
				 }
		    }
        selectWindow("Results"); 
   		run("Close"); 
        selectWindow(filename);
		roiManager("Show None");
		roiManager("Show All");

		roiManager("measure");
		
		if (X>=100)  {
			saveAs("Results", Results_file + "Point_Fluo_"+ X +".csv");
			}
		else
			if (X>=10)  {
			saveAs("Results", Results_file + "Point_Fluo_0"+ X +".csv");
			}
			else {
				saveAs("Results", Results_file + "Point_Fluo_00"+ X +".csv");
				}
		selectWindow(Mask_filename);
		close ();
        selectWindow(filename);
        close ();

        selectWindow("Results"); 
   		run("Close"); 
		roiManager("save", ROI_file + filename + ".zip");
   		roiManager("reset");
		}
```
