## Supplementary material for "Dynamics of drug response in single mycobacterial cells by microfluidic dose-response assay": Data S1-S7: Ch-Extraction.ijm

```
//--------------------------------------------------------------------------------------------------------------------
// DATA SOURCES
//--------------------------------------------------------------------------------------------------------------------


ExpFile = "/Users/mmistret/Documents/Data/Microfluidic/Longterme_Imaging/05_Gradiant_Chip/03_5C_MTB/02_MOX/20210422_5C_MTB_250_MOX_R2/"


//--------------------------------------------------------------------------------------------------------------------
//--------------------------------------------------------------------------------------------------------------------

//Generare all folder for saving data
//Folder for BrightField
File.makeDirectory(ExpFile + "BF_TIFF");

//Folder for GFP
File.makeDirectory(ExpFile + "GFP_TIFF");

//Folder for Mcherry
File.makeDirectory(ExpFile + "DSRed_TIFF");

//Folder for Mask V1 fluorecence based
File.makeDirectory(ExpFile + "Mask_V1");

//Folder for Results 
File.makeDirectory(ExpFile + "Results");

//Folder for ROI
File.makeDirectory(ExpFile + "ROI");


//--------------------------------------------------------------------------------------------------------------------

//DV File
DV_file = 		ExpFile + "DV/";
//BF_Channel_TIFF
BF_image = 		ExpFile + "BF_TIFF/";
//FITC_Channel_TIFF
GFP_image =		 ExpFile + "GFP_TIFF/";
//FITC_Channel_TIFF
DSRed_image = 	ExpFile + "DSRed_TIFF/";
//--------------------------------------------------------------------------------------------------------------------

setBatchMode(true); 
list = getFileList(DV_file);

        X=1;	
        print(DV_file);

for (i = 0; i <list.length; i++) {
        print(list[i]);
        //ExtractBF(DV_file, BF_image, list[i], X);
       ExtractGFP(DV_file, GFP_image, list[i], X);
      ExtractDSRed(DV_file, DSRed_image, list[i], X);
		X = X + 1;
		}
setBatchMode(false)

//--------------------------------------------------------------------------------------------------------------------
//BF Channel extraction
//--------------------------------------------------------------------------------------------------------------------
function ExtractBF(DV_file, BF_image, filename, X) {

		run("Bio-Formats", "open="+ DV_file + filename +" autoscale color_mode=Default view=[Standard ImageJ] stack_order=Default");


        getDimensions(width, height, channels, slices, frames);

        run("Stack to Hyperstack...", "order=xyczt(default) channels="+ channels +" slices=1 frames="+ frames +" display=Color");
		run("Next Slice [>]");
		resetMinAndMax();
		
		run("Reduce Dimensionality...", "frames");  

		if (X>=100)  {
			saveAs("TIFF", BF_image + "Point_BF_"+ X );
			}
		else
			if (X>=10)  {
			saveAs("TIFF",  BF_image + "Point_BF_0"+ X );
			}
			else {
				saveAs("TIFF",  BF_image + "Point_BF_00"+ X );
				}
		
		close ();
		}
		
//--------------------------------------------------------------------------------------------------------------------
//GFP Channel extraction
//--------------------------------------------------------------------------------------------------------------------
function ExtractGFP(DV_file, GFP_image, filename, X) {


		run("Bio-Formats", "open="+ DV_file + filename +" autoscale color_mode=Default view=[Standard ImageJ] stack_order=Default");

        getDimensions(width, height, channels, slices, frames);

	    
       run("Stack to Hyperstack...", "order=xyczt(default) channels="+ channels+" slices=1 frames="+ frames +" display=Color");
		run("Next Slice [>]");
		run("Next Slice [>]");
		run("Next Slice [>]");

		
		resetMinAndMax();	
		
		run("Reduce Dimensionality...", "frames");  
		//run("Subtract Background...", "rolling=50 stack");

		if (X>=100)  {
			saveAs("TIFF", GFP_image + "Point_GFP_"+ X );
			}
		else
			if (X>=10)  {
			saveAs("TIFF", GFP_image + "Point_GFP_0"+ X );
			}
			else {
				saveAs("TIFF", GFP_image + "Point_GFP_00"+ X );
				}
		
		close ();
		}

//--------------------------------------------------------------------------------------------------------------------
//--------------------------------------------------------------------------------------------------------------------
//Mcherry Channel extraction
//--------------------------------------------------------------------------------------------------------------------
function ExtractDSRed(DV_file, DSRed_image, filename, X) {


		run("Bio-Formats", "open="+ DV_file + filename +" autoscale color_mode=Default view=[Standard ImageJ] stack_order=Default");
        getDimensions(width, height, channels, slices, frames);

	    
       	run("Stack to Hyperstack...", "order=xyczt(default) channels="+ channels+" slices=1 frames="+ frames +" display=Color");
		run("Next Slice [>]");
		run("Next Slice [>]");


		run("Reduce Dimensionality...", "frames");   
		//run("Subtract Background...", "rolling=50 stack");
		resetMinAndMax();	

		if (X>=100)  {
			saveAs("TIFF", DSRed_image + "Point_DSRed_"+ X );
			}
		else
			if (X>=10)  {
			saveAs("TIFF", DSRed_image + "Point_DSRed_0"+ X );
			}
			else {
				saveAs("TIFF", DSRed_image + "Point_DSRed_00"+ X );
				}
		
		close ();
		}
```
