## Supplementary material for "Dynamics of drug response in single mycobacterial cells by microfluidic dose-response assay": Data S1-S7: csv-Management.R

```
################################################################
##                                                            ##
##     CVF MANAGEMENT                                         ##
##                                                            ##
##     Gael A. Millot                                         ##
##     Version v1.0.0                                         ##
##                                                            ##
################################################################


################################ Introduction


# Compatible with R v3.5.1
# The code in the "Parameters that need to be set by the user" section indicates the code that has to be personalized
# Restrictions and hypotheses:
# 1) the folder must only contain files to process
# 2) In file names, the character string between the 2nd and 3rd "_" must be the file number
# 3) files must contain the following columns names: "Labels", "Area", "Mean", "StdDev"
# 4) Label column must contain names with caracter string after the last ":" character being the frame number


################################ End Introduction


################################ Initialization


# R version checking
if(version$version.string != "R version 3.5.1 (2018-07-02)"){
    cat("\n\nBEWARE: THE ", version$version.string, " IS NOT THE 3.5.1 RECOMMANDED\n\n")
}
# Packages checking
if( ! c("lubridate") %in% rownames(installed.packages(lib.loc = .libPaths()))){
stop(paste0("\n\n================\n\nERROR: PACKAGE lubridate MUST BE INSTALLED IN AT LEAST ONE OF THESE DIRECTORIES:\n", paste(.libPaths(), collapse = "\n"), "\n\n================\n\n"))
}
# other initializations
erase.objects = TRUE # write TRUE to erase all the existing objects in R before starting the algorithm and FALSE otherwise. Beginners should use TRUE
if(erase.objects == TRUE){
    rm(list=ls())
    erase.objects = TRUE
}
erase.graphs = FALSE # write TRUE to erase all the graphic windows in R before starting the algorithm and FALSE otherwise


################################ End Initialization


################################ Parameters that need to be set by the user


################ File names and locations


project.name <- "Mistretta_pro12193_CSV_management"
path.in <-  "/Users/mmistret/Documents/Data/Microfluidic/Longterme_Imaging/05_Gradiant_Chip/02_5C_MSmeg/20201127_5C_MS_M02_MOX_R2/EXP1/Results_GFP/" # Define the absolute pathway of the folder containing the input data files
path.out <- "/Users/mmistret/Documents/Data/Microfluidic/Longterme_Imaging/05_Gradiant_Chip/02_5C_MSmeg/20201127_5C_MS_M02_MOX_R2/EXP1/" # Define the absolute pathway of the destination folder that will contain the results (exported data)
path.function1 <- "/Users/mmistret/Documents/Software/HubTool/cute_little_functions-v2.0.0/" # Define the absolute pathway of the folder containing functions created by Gael Millot. See https://gitlab.pasteur.fr/gmillot/cute_little_functions
name.source.file1 <- "cute_little_functions.R"


################ Parameters


colum.name.recovery <- c("Area", "Mean", "StdDev") # column names that must be present in files from path.in
col.sep <- "," # separator of columns inside files. Write "\t" for tabulations
optional.text <- "" # write here an optional text to include in results and graphs


################################ End Parameters that need to be set by the user


################################ Recording of the initial parameters


param.list <- c(
    "erase.objects", 
    "erase.graphs", 
    "project.name", 
    "path.in", 
    "path.out", 
    "path.function1", 
    "name.source.file1", 
    "colum.name.recovery",
    "col.sep",
    "optional.text"
)
if(any(duplicated(param.list))){
    stop(paste0("\n\n================\n\nERROR: THE param.list OBJECT CONTAINS DUPLICATED ELEMENTS\n\n================\n\n")) # message for developers
}
if(erase.objects == TRUE){
    created.object.control <- ls()[ ! ls() %in% "param.list"]
    if( ! (all(created.object.control %in% param.list) & all(param.list %in% created.object.control))){
        stop(paste0("\n\n================\n\nERROR: INCONSISTENCIES BETWEEN THE param.list ELEMENTS AND THE CREATED OBJECTS\nTHE CREATED OBJECTS (created.object.control) NOT PRESENT IN THE param.list ARE: ", paste(created.object.control[ ! created.object.control %in% param.list], collapse = " "), "\nTHE param.list ELEMENTS NOT PRESENT IN THE CREATED OBJECTS (created.object.control) ARE: ", paste(param.list[ ! param.list %in% created.object.control], collapse = " "), "\n\n================\n\n")) # message for developers
    }
}
char.length <- nchar(param.list)
space.add <- max(char.length) - char.length + 5
param.ini.settings <- NULL
for(i in 1:length(param.list)){
    param.ini.settings <- c(param.ini.settings, paste0("\n", param.list[i], paste0(rep(" ", space.add[i]), collapse = ""), paste0(get(param.list[i]), collapse = ",")))
}


################################ End Recording of the initial parameters


################################ Main code


################ Pre-ignition checking


# settings
check.ok <- TRUE
tempo.check.script <- '
if(tempo$problem == TRUE){
check.ok <- FALSE
}
'
# import functions
if( ! file.exists(paste0(path.function1, name.source.file1))){
    tempo.cat <- paste0("\n\n================\n\nERROR: FILE NAME GIVEN BY path.function1 AND name.source.file1 PARAMETERS DOES NOT EXISTS: ", paste0(path.function1, name.source.file1), "\n\n================\n\n")
    cat(tempo.cat)
    check.ok <- FALSE
}else{
    source(paste0(path.function1, name.source.file1)) # call the "fun_param_check", "fun_export_data"
}
# file name and locations
tempo <- fun_param_check(data = project.name, class = "character", length = 1) ; eval(parse(text = tempo.check.script)) # execute tempo.script
tempo <- fun_param_check(data = path.in, class = "character", length = 1) ; eval(parse(text = tempo.check.script)) # execute tempo.script
tempo <- fun_param_check(data = path.out, class = "character", length = 1) ; eval(parse(text = tempo.check.script)) # execute tempo.script
tempo <- fun_param_check(data = path.function1, class = "character", length = 1) ; eval(parse(text = tempo.check.script)) # execute tempo.script
tempo <- fun_param_check(data = name.source.file1, class = "character", length = 1) ; eval(parse(text = tempo.check.script)) # execute tempo.script
# parameters
tempo <- fun_param_check(data = colum.name.recovery, class = "character") ; eval(parse(text = tempo.check.script)) # execute tempo.script
tempo <- fun_param_check(data = col.sep, class = "character", length = 1) ; eval(parse(text = tempo.check.script)) # execute tempo.script
tempo <- fun_param_check(data = optional.text, class = "character", length = 1) ; eval(parse(text = tempo.check.script)) # execute tempo.script


################ Ignition


options(scipen = 7) #affichage Scientific ou std
 check.ok <- TRUE
ini.time <- as.numeric(Sys.time()) # time of process begin, converted into seconds
analysis.nb <- trunc(ini.time) # to provide a specific number ot each analysis
name.dir <- paste0(project.name, "_", analysis.nb)
if( ! dir.exists(path.out)){
    tempo.cat <- paste0("\n\n============\n\nERROR: FILE DIRECTORY INDICATED IN THE path.out PARAMETER DOES NOT EXISTS: ", path.out, "\n\n============\n\n")
    cat(tempo.cat)
    check.ok <- FALSE
}else{
    path.out <- paste0(path.out, name.dir)
    suppressWarnings(dir.create(path.out))
}
log.file <- paste0("log_", analysis.nb,".txt")
fun_export_data(data = paste0("\n\n################################ ", project.name, "_", analysis.nb, " ################"), output = log.file, no.overwrite = FALSE, path = path.out, sep = 4)


################ Pre-import checking


if( ! dir.exists(path.in)){
    tempo.cat <- paste0("\n\n============\n\nERROR: FILE DIRECTORY INDICATED IN THE path.in PARAMETER DOES NOT EXISTS: ", path.out, "\n\n============\n\n")
    cat(tempo.cat)
    check.ok <- FALSE
}

if(length(list.files(path.in)) == 0){
    tempo.cat <- paste0("\n\n============\n\nERROR: FILE DIRECTORY INDICATED IN THE path.in PARAMETER IS EMPTY: ", path.in, "\n\n============\n\n")
    cat(tempo.cat)
    check.ok <- FALSE
}else{
    file.list <- list.files(path.in)
    file.list.split <- strsplit(file.list, split = "\\.") # name split according to dot in file names
    if(length(table(sapply(file.list.split, length))) != 1){
        tempo.cat <- paste0("\n\n============\n\nERROR: FILES NAMES IN ", path.in, " DO NOT HAVE THE SAME STRUCTURE (DIFFERENT NUMBER OF DOT CHARACTER \".\" IN NAMES)\n\n============\n\n")
        cat(tempo.cat)
        check.ok <- FALSE
    }else if(length(table(sapply(file.list.split, "[[", length(file.list.split[[1]])))) != 1){
        tempo.cat <- paste0("\n\n============\n\nERROR: FILES NAMES IN ", path.in, " DO NOT HAVE THE SAME EXTENSION (AFTER LAST \".\" IN NAMES): ", paste(unique(sapply(file.list.split, "[[", length(file.list.split[[1]]))), collapse = " "), "\n\n============\n\n")
        cat(tempo.cat)
        check.ok <- FALSE
    }
}


################ Code after checking


if(check.ok == TRUE){
    # Checking similar internal file structure
    max.nrow <- 0
    col.name <- NULL
    for(i2 in 1:length(file.list)){
        ini.user.data <- read.table(paste0(path.in, file.list[i2]), sep = col.sep, header = TRUE, comment.char = "", stringsAsFactors = FALSE) # Data import
        if(! all(colum.name.recovery %in% names(ini.user.data))){
            tempo.cat <- paste0("\n\n============\n\nERROR: COLUMN NAMES ", paste(colum.name.recovery, collapse = " "), " INDICATED IN THE colum.name.recovery PARAMETER ARE MISSING IN THE FOLLOWING FILE", paste0(path.in, file.list[i2]), "\n\n============\n\n")
            fun_export_data(path = path.out, data = tempo.cat, output = log.file)
            stop(tempo.cat)
        }else{
            col.name <- c(col.name, paste0("file.", strsplit(file.list[i2], split = "_")[[1]][3])) #!!!!!Take third underscore where is the number
            if(nrow(ini.user.data) > max.nrow){
                max.nrow <- nrow(ini.user.data)
            }
        }
    }
    col.name <- sort(col.name)
    # Data recovering
    for(i2 in 1:length(colum.name.recovery)){
        assign(colum.name.recovery[i2], as.data.frame(matrix(NA, nrow = max.nrow, ncol = length(file.list), dimnames = list(paste0("frame.", 1:max.nrow), col.name)), header = TRUE, stringsAsFactors = FALSE))
    }
    for(i2 in 1:length(file.list)){
        ini.user.data <- read.table(paste0(path.in, file.list[i2]), sep = col.sep, header = TRUE, comment.char = "", stringsAsFactors = FALSE) # Data import
        tempo.frame.num.list <- strsplit(ini.user.data[, "Label"], split= ":") #!!!! Need 3 ":" and take frame after the third one
        if(length(table(sapply(tempo.frame.num.list, length))) != 1){    
            tempo.cat <- paste0("\n\n============\n\nERROR: NAMES IN COLUMN ", "Label", " OF FILE ", paste0(path.in, file.list[i2]), " DO NOT HAVE THE SAME STRUCTURE (DIFFERENT NUMBER OF COLON CHARACTER \":\" IN NAMES)\n\n============\n\n")
            fun_export_data(path = path.out, data = tempo.cat, output = log.file)
            stop(tempo.cat)
        }else{
            tempo.frame.num <- as.numeric(sapply(tempo.frame.num.list, "[[", length(tempo.frame.num.list[[1]]))) # recover the string after last ":"
            tempo.file.num <- paste0("file.", strsplit(file.list[i2], split = "_")[[1]][3])
            for(i3 in 1:length(colum.name.recovery)){
                assign(colum.name.recovery[i3], {x <- get(colum.name.recovery[i3]) ; x[tempo.frame.num, tempo.file.num] <- ini.user.data[, colum.name.recovery[i3]]; x})
            }
        }
    }
    for(i2 in 1:length(colum.name.recovery)){
        write.table(get(colum.name.recovery[i2]), file = paste0(path.out, "/", colum.name.recovery[i2], ".txt"), row.names = TRUE, col.names = NA, append = FALSE, quote = FALSE, sep = "\t") # col.names = NA to insert an empty tabulation for row names in the header (see ?write.table)
        write.csv(get(colum.name.recovery[i2]), file = paste0(path.out, "/", colum.name.recovery[i2], ".csv"))
        write.csv2(get(colum.name.recovery[i2]), file = paste0(path.out, "/", colum.name.recovery[i2], ".csv2"))
    }


############ total computing time
    
    
    fin.time <- as.numeric(Sys.time()) # time of process end
    suppressPackageStartupMessages(library(lubridate, quietly = TRUE)) # provide the seconds_to_period() function, qui whcih converts seconds in days, hours, minutes, seconds
    cat("TOTAL COMPUTATION TIME: ", as.character(seconds_to_period(round(fin.time - ini.time))), "\n\n")
    fun_export_data(path = path.out, data = paste0("\n\n\n\n################################ TOTAL COMPUTING TIME\n\n", as.character(seconds_to_period(round(fin.time - ini.time)))), output = log.file, sep = 4)
    
    
############ Parameter printing


}else if(check.ok == FALSE){
    tempo.cat <- paste0("\n\n============\n\nERROR: PROBLEM DETECTED WITH check.ok, \n\n============\n\n")
    cat(tempo.cat)
}
fun_export_data("\n\n\n\n################################ INITIAL SETTINGS OF PARAMETERS", output = log.file, path = path.out)
fun_export_data(path = path.out, data = param.ini.settings, output = log.file, vector.cat = TRUE)


################################ End Main code
```
