## Supplementary material for "Dynamics of drug response in single mycobacterial cells by microfluidic dose-response assay": Data S1-S7: GFP-Mask.ijm

```
//Fluo_Channel Image
input = "/Users/mmistret/Documents/Data/Microfluidic/Longterme_Imaging/05_Gradiant_Chip/03_5C_MTB/20200203_5C_MTB_MOX_250_R1/02_Image/GFP_TIFF/";
//Mask file
output = "/Users/mmistret/Documents/Data/Microfluidic/Longterme_Imaging/05_Gradiant_Chip/03_5C_MTB/20200203_5C_MTB_MOX_250_R1/02_Image/Mask_GFP/";

setBatchMode(true); 
list = getFileList(input);
X=1;
for (i = 0; i <list.length; i++) {
        Mask1(input, output, list[i], X);
        X = X+1 ; }
setBatchMode(false)


function Mask1(input, output, filename, X) {
        open(input + filename);
        
		run("Gaussian Blur...", "sigma=2 stack");
		run("Smooth", "stack");
		run("Smooth", "stack");
		run("Smooth", "stack");
		run("Smooth", "stack");
		run("Smooth", "stack");
		run("Smooth", "stack");

		resetMinAndMax();
		setSlice(30);
		
		setOption("BlackBackground", true);
		run("Convert to Mask", "method=Intermodes background=Default calculate black");
		run("Erode", "stack");
		run("Dilate", "stack");
		
	
		if (X>=100)  {
			saveAs("TIFF", output + "Mask_GFP_V1_"+ X );
			}
		else
			if (X>=10)  {
			saveAs("TIFF", output + "Mask_GFP_V1_0"+ X );
			}
			else {
				saveAs("TIFF", output + "Mask_GFP_V1_00"+ X );
				}
		close();
        }
```
